# Rapid Homolog Juxtaposition During Meiotic Chromosome Pairing

**DOI:** 10.1101/2024.03.23.586418

**Authors:** Tadasu Nozaki, Beth Weiner, Nancy Kleckner

## Abstract

A central basic feature of meiosis is pairing of homologous maternal and paternal chromosomes (“homologs”) intimately along their lengths. Recognition between homologs and their juxtaposition in space are mediated by axis-associated DNA recombination complexes. Additional effects ensure that pairing occurs without ultimately giving entanglements among unrelated chromosomes. Here we examine the process of homolog juxtaposition in real time by 4D fluorescence imaging of tagged chromosomal loci at high spatio-temporal resolution in budding yeast. We discover that corresponding loci start coming together from a quite large distance (∼1.8 µm) and progress to completion of pairing in a very short time, usually less than six minutes (thus, “rapid homolog juxtaposition” or “RHJ”). Juxtaposition initiates by motion-mediated extension of a nascent interhomolog DNA linkage, raising the possibility of a tension-mediated trigger. In a first transition, homolog loci move rapidly together (in ∼30 sec, at speeds of up to ∼60 nm/sec) into a discrete intermediate state corresponding to canonical ∼400 nm axis distance coalignment. Then, after a short pause, crossover/noncrossover differentiation (crossover interference) mediates a second short, rapid transition that brings homologs even closer together. If synaptonemal complex (SC) component Zip1 is present, this transition concomitantly gives final close pairing by axis juxtaposition at ∼100 nm, the “SC distance”. We also find that: (i) RHJ occurs after chromosomes acquire their prophase chromosome organization; (ii) is nearly synchronously over thirds (or more) of chromosome lengths; but (iii) is asynchronous throughout the genome. Furthermore, cytoskeleton-mediated movement is important for the timing and distance of RHJ onset and also for ensuring normal progression. Potential implications for local and global aspects of pairing are discussed.

## Introduction

Meiosis is the modified cellular program that underlies gamete formation for sexual reproduction and shuffling of genetic information as the substrate for evolution. A central unique feature of this program is pairing of maternal and paternal chromosomes (“homologs”) along their lengths, which occurs in temporal and functional linkage with programmed recombination during a prolonged prophase period ^1^. In the present study we sought to elucidate the nature of homolog juxtaposition by monitoring the positions of individual tagged loci, in 3D in real time (4D), at high spatio-temporal resolution, in budding yeast. We reasoned that this approach, being unique as compared to previous approaches, should provide new perspectives, orthogonal and complementary to those provided by previous studies.

The existence of homolog pairing was first provided by classical light microscopy, which revealed very tight association (“pachytene synapsis”) and showed that this final state was preceded by a tendency for association at greater distances. Subsequent electron microscope (EM) and fluorescence imaging analysis revealed that synapsis corresponds to formation of the synaptonemal complex (“SC”), a universal structure, in which close-packed transverse filaments links paired axes at a distance of ∼100 nm (Fig. 1a right) ^1^. Furthermore, in *Sordaria macrospora*, the preceding distance association has been defined to specifically comprise coalignment of homolog axes at ∼400 nm (Fig. 1a left) ^1^. In addition, the two stages are linked by emergence of cytologically-prominent inter-axis bridges, at least some of which are sites of SC nucleation (Fig. 1a middle) ^1^.

**Fig. 1.**
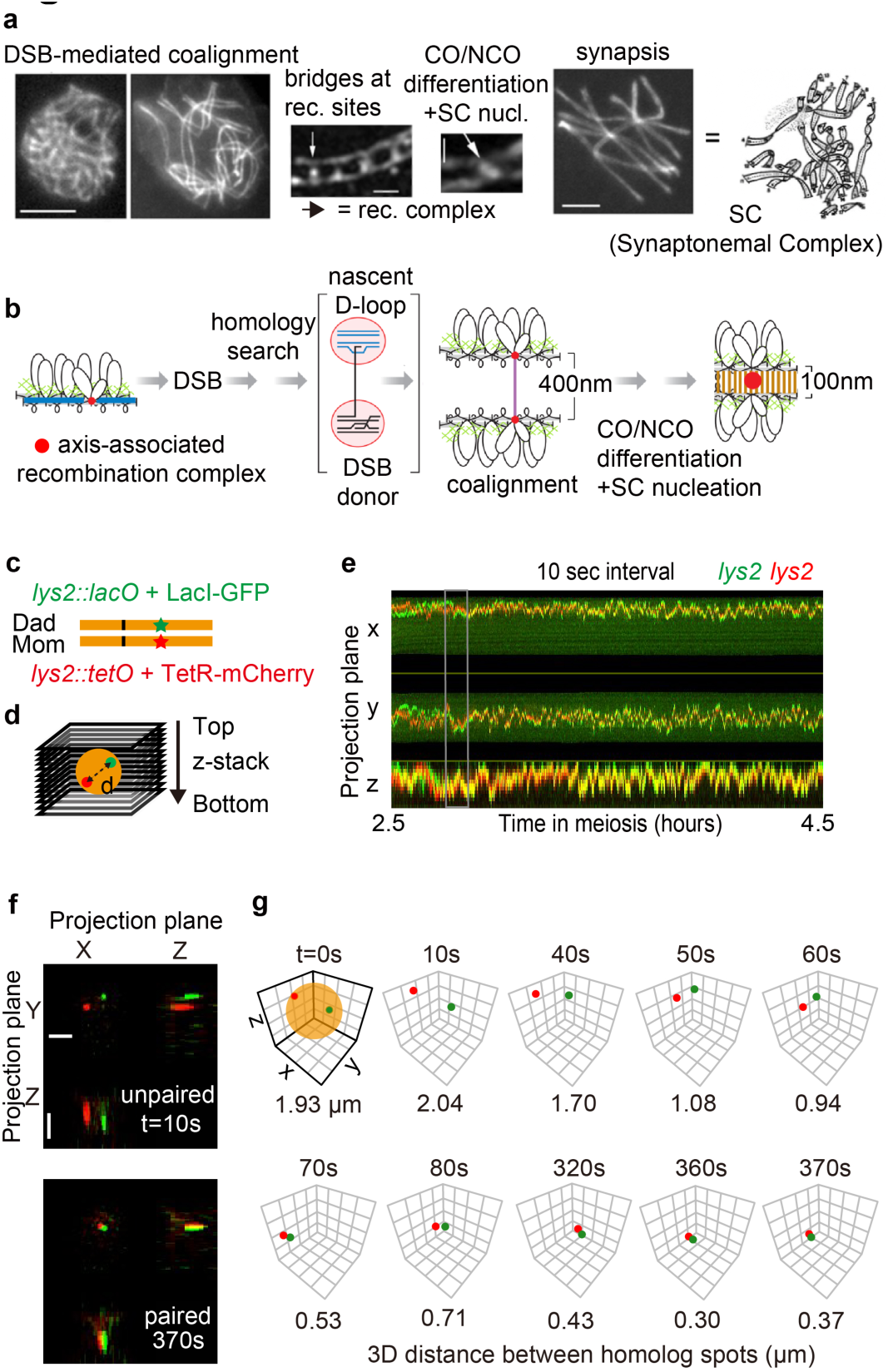
Homolog pairing by progressive recombination-mediated juxtaposition. **a.** Juxtaposition of homolog axes as illustrated by *Sordaria macrospora* ^1,19^. DSB-initiated recombination mediates coalignment of axes to 400 nm; DNA/structure bridges then emerge at sites of recombination interactions and are the substrate for interference-mediated patterning of crossovers (CO) and concomitant installation of the synaptonemal complex (SC), which juxtaposes axes to ∼100 nm. Scale bars in right and left figures = 2 µm. Scale bars in middle figures = 200 nm. **b.** Axis juxtaposition is mediated by recombination events involving nascent D-loop(s) in axis-associated complexes ^1^. **c.** Pairing was assayed by imaging of *lacO*/LacI-mEGFP and *tetO*/TetR-mCherry tags located at corresponding positions on homologs, e.g. at the *LYS2* locus. **d.** Each image of a 3D time-lapse comprises a two-color Z-stack that allows to define the positions of the two fluorescent spots and their 3D distance (“d”); orange indicates approximate nucleus size. **e.** Kymograph of 3D spot images in the three projection planes for *lys2*-green and *lys2*-red (panel **c**) at 10-sec intervals for 2 hours. **f, g.** Illustration of progression of homolog pairing from far apart (unpaired) to close together (paired) during the indicated time period of panel **e** (gray lines). Representative 3D image of two-colored spots projected onto XY, XZ and YZ planes. Top, white scale bars = 2 µm (**f**). 3D plots and 3D inter-spot distances are shown. *x*, *y*, *z*-axes = 4 µm; orange indicates approximate nucleus size, ∼2 µm diameter (**g**).

It is now also known that these global outcomes of distance coalignment and SC formation are mediated by the local events of DNA recombination (Fig. 1b). Meiotic recombination initiates by programmed double-strand breaks (DSBs). The first contact between homologs involves a nascent D-loop between the single-strand DNA “tail” at one DSB end and the corresponding region on the DNA of a non-sister (homolog) chromatid ^2^. Exactly when and how this association occurs, and the steps leading to a ∼400 nm coalignment interaction, are unknown. As a general principle, however, events at the DNA level can result in juxtaposition of homologs at the whole chromosome level because the biochemical complexes of recombination are physically and functionally associated with their underlying structural axes, with local effects transmitted along the chromosomes via these structures ^1^. The transition from coalignment to SC formation is also mediated by recombination/structure linkage. SC installation nucleates at sites of recombination, as seen in several organisms ^1^. In *Sordaria macrospora,* where relevant events are well defined, this transition is also known to be triggered by crossover/noncrossover differentiation, as governed by the classical phenomenon of crossover interference, which concomitantly results in closer juxtaposition of axes and SC nucleation (Fig. 1a, b; below) ^3–7^.

Importantly, the events of recombination-mediated homolog juxtaposition are part of a global program in which large numbers of unrelated chromosomes, often long and thin and closely-packed, manage to sort themselves into topologically regular pairs. Multiple components have been shown, or proposed, to work collaboratively to promote the avoidance, minimization and/or resolution of unacceptable entanglements ^1^. How local juxtaposition at individual recombination sites is integrated with these global features is unclear.

Another central feature of meiosis that contributes to all of these effects is cytoskeleton-mediated end-led motion ^8–14^. Chromosome telomeres are specifically linked to, and then through, the nuclear envelope via “LINC” complexes which, in turn, are associated with motor proteins that move along cytoskeletal actin or tubulin filaments. This end-led motion is variously proposed to promote juxtaposition of homologs, to serve as a stringency factor to eliminate unwanted linkages, and/or to actively move entrapped chromosomes out of their “interlocked” state ^1,9,11^.

The present study further addresses the complexities of meiotic homolog pairing by examining the spatial juxtaposition of individual homolog loci in real-time. Corresponding positions on homologs of budding yeast chromosomes were marked with fluorescent tags and the relative positions of the corresponding “spots” were defined in 4D (3D over time) at high spatio-temporal resolution over long time periods. This approach reveals that corresponding loci on homologs undergo rapid directed juxtaposition that initiates from a significant (potentially conserved) distance and is triggered by a discrete increase in that distance. We propose that onset of juxtaposition is triggered by tension along the nascent interhomolog connection. Homolog loci first come together rapidly to discrete intermediate stage corresponding to canonical distance coalignment. The mechanistic basis for this effect now remains to be defined. Finally, after a short pause, crossover/noncrossover differentiation promotes closer pairing and concomitant SC formation. Most notably, this entire series of events, from onset to close pairing/SC formation, occurs in just a few minutes (hence “Rapid Homolog Juxtaposition” (RHJ)). Additionally, RHJ onset occurs nearly synchronously at nearby loci on the same chromosome. Finally, new roles for end-led motion are revealed, for both onset and progression of RHJ. These findings have several implications for global nucleus-wide aspects of homolog pairing. Together they provide a new foundation for further analysis of homolog coalignment, crossover/noncrossover differentiation, end-led motion and their relationships within, along and among chromosomes.

## Results

### Experimental Approach

Budding yeast chromosomal loci of interest were tagged with arrays of *lac* or *tet* operators and visualized using cognate repressors tagged with mEGFP or mCherry, respectively (Fig. 1c). Cell cultures were taken through synchronous meiosis (Materials and Methods) and fluorescent “spots” in individual nuclei were visualized over time by wide-field 3D fluorescence imaging. Images were analyzed using a specially-designed low signal-to-noise ratio (SNR) algorithm and imaging setup (Extended Data Fig. 1a) ^12^. 3D images (Fig. 1d) were usually acquired at intervals of ten seconds for 2 hours (e.g. Fig. 1e, f). In some cases, images were obtained at one-minute intervals for ≥ 8 hours, allowing definition of homolog spot positions from earliest meiosis through both meiotic divisions (Extended Data Fig. 1b-g, Supplementary Video 1). A representative time series illustrates our main finding: homolog spots move from wide separation to close juxtaposition very rapidly, usually in less than six minutes (Fig. 1g, Supplementary Video 2).

The dynamics of locus-specific juxtaposition was analyzed for six different individual locus pairs located on three different chromosomes of varying lengths (n=61 cells; Extended Data Fig. 2a). Findings are often illustrated for the *lys2* locus (n=14 cells), which is located internally on one of yeast’s longer chromosomes (ChrII), far from the centromere and telomeres (Fig. 1c). The same basic features are seen at all analyzed loci (below; Extended Data Fig. 2b, Supplementary Video 3).

### Multi-step Rapid Homolog Juxtaposition (RHJ)

Tagged homolog loci become juxtaposed in three basic stages, clear by visual inspection, and defined precisely by a step-detection algorithm (Fig. 2a-c, Extended Data Fig. 2b and 3). As illustrated for *lys2*: (i) Inter-focus distances initially fluctuate around a mean of ∼1.4 µm for many minutes (cyan). (ii) At a certain point, homolog spots undergo a rapid decrease of ∼600 nm in inter-focus distance, to ∼0.8 µm, which comprises a discrete intermediate stage (orange). (iii) After a short pause, inter-focus distance again rapidly decreases by ∼340 nm to a close pairing distance of ∼0.45 µm (purple). Close pairing is stable until onset of the MI division (Extended Data Fig. 1c and f). In rare exceptional cases, the intermediate stage is not detectable, an extra stage is prominent before the intermediate stage, or close pairing is temporarily unstable (Extended Data Fig. 2 c-e).

**Fig. 2.**
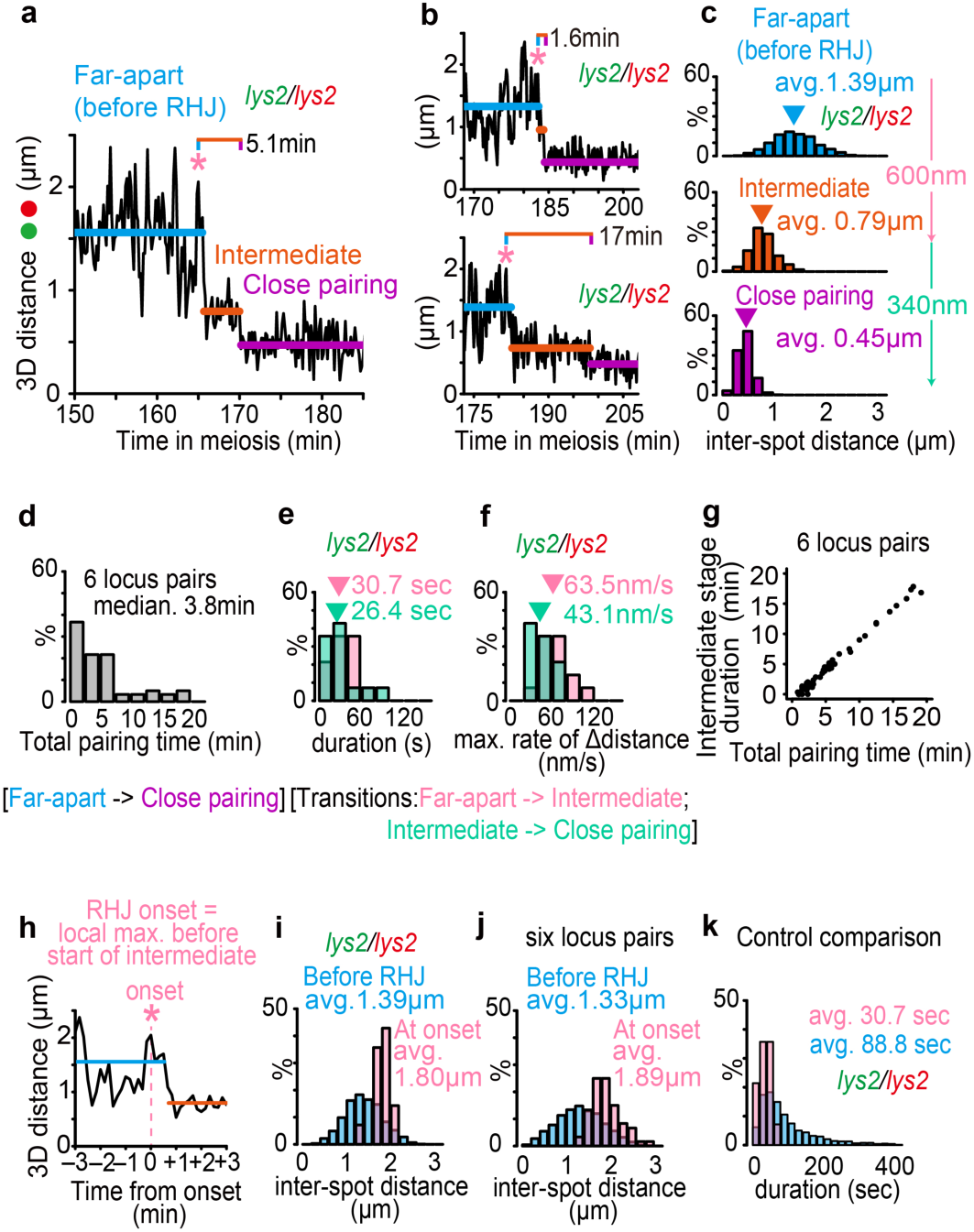
Rapid homolog juxtaposition (RHJ). **a-g.** Three stages of RHJ. **a, b.** Representative examples of 3D distances between tagged homolog loci (*lys2*) over time at 10-second intervals. Three states, apparent by eyes, are defined rigorously by a step-detection function (Extended Data Fig. 3a, Materials and Methods): far apart (before RHJ), cyan; intermediate, orange; and close pairing, purple. Total pairing time from RHJ onset (asterisk; see panel **h** below) to close pairing indicated (see the details in Extended Data Fig. 3b). **c.** Distribution of inter-spot distances during each of the three stages in (**a**, **b**) (*lys2*; n=14 cells). **d.** Distribution of total pairing times as defined in panels **a, b** at six different locus pairs (n=60 nuclei; Extended Data Fig. 2a). **e, f.** Two RHJ transitions are both short and rapid as shown by their durations (**e**) and maximum 10 sec interval speeds (**f**) (*lys2*; n = 14 cells). **g.** The intermediate stage (ordinate) accounts for almost all of the total pairing time (abscissa; from **d**) (n=60, 6 locus pairs). **h-k.** RHJ is initiated by a specific increase in inter-homolog distance to ∼1.8 µm. **h.** Onset of RHJ is defined as the local maximum of inter-spot distance that immediately proceeds progression to the intermediate stage (asterisk) (Extended Data Fig. 3b). **i, j.** The inter-spot distance at RHJ onset as defined in panel **h** (pink) is significantly larger than the inter-spot distances at all preceding time points (cyan) at *lys2* (**i**; n=14 cells); and six locus pairs (**j**; n=60 cells). **k.** Control showing that the “onset” peak in panel **h** is specifically, temporally linked to occurrence of RHJ. The time between onset and achieving the intermediate stage is a narrow peak with an average of ∼30 sec (pink) (from panel **e**). In contrast, prior to onset, for all time points at which the inter-spot distance is in the same range as the intermediate stage (>0.6 µm and <1.2 µm), the analogous preceding local maximum are more broadly distributed and significantly longer on average (∼90 sec) (cyan, n=14 cells, see the details in Extended Data Fig. 6c, d).

Remarkably, progression from “far apart” to “close pairing” usually occurs in just a few minutes, thus defining the existence of “rapid homolog juxtaposition” (RHJ). The median duration for all analyzed nuclei was 3.8 minutes (Fig. 2d, n=60 cells). Median duration was 5.8 min at *lys2* and 2.1-5.5 min at other loci (Extended Data Fig. 4a).

The RHJ transitions into and beyond the intermediate stage, both, are very short (∼30 sec; Fig. 2e). Both also involve very rapid coming-together of homolog spots, with maximum speeds of ∼60 and 40 nm/sec as defined by corresponding 10 sec intervals (Fig. 2f). These rates are faster than fluctuations in the distance between loci prior to the onset of RHJ, implying that both transitions involve active, directed processes (Extended Data Fig. 5; Discussion). Given that the two transitions are very short, the total duration of RHJ is essentially determined by the length of the intermediate stage, which ranges from 0 to 17.8 min (median 2.8 min) in direct correlation with total pairing time (Fig. 2g, n=60 cells).

### Onset of RHJ is initiated by a discrete increase in inter-spot distance

Prior to RHJ, inter-spot distances fluctuate dramatically (Fig. 2a and b, Extended Data Fig. 2b) due to cytoskeleton-mediated end-led chromosome motion (below). Within this fluctuating landscape, RHJ is always preceded by a prominent local peak of inter-spot distance, seen as a preceding local maximum in inter-spot distance (asterisk, Fig. 2h). At *lys2,* inter-spot distances at this RHJ “onset peak” are narrowly distributed around a value at average 1.80 µm, significantly higher than the pre-RHJ average of 1.39 µm (Fig. 2i; pink versus cyan, *t*-test *P* = 1.4 × 10^-6^, n=14 cells, Extended Data Fig. 6a). Similar onset distances are observed at all loci (Fig. 2j, average 1.89 µm for onset peak and average 1.33 µm for pre-RHJ, *t*-test *P* < 2.2 × 10^-16^, n=60 cells, Extended Data Fig. 4b and 6b).

Control analysis confirms that this immediately preceding local maximum is temporally correlated with (and thus functionally correlated with) the onset of RHJ. During RHJ, the time between the occurrence of the onset peak and the time at which the intermediate stage is achieved is narrowly distributed around ∼30 sec on average (range 10-70 sec) (Fig. 2k pink). As a control comparison, we identified all time points prior to RHJ at which homolog spots fortuitously occurred at the same distance as at the true intermediate stage (range of 0.6 – 1.2 µm) and, in each case, identified the corresponding immediately preceding local maximum of inter-spot distance. The time between the occurrence of these two functionally uncorrelated events is 2-3 times longer (∼90 sec on average), and much more broadly distributed, than those for RHJ-associated events (Fig. 2k cyan, Kolmogorov-Smirnov test *P* = 1.5 × 10^-3^, Extended Data Fig. 6c, d).

An analogous discrete extension may also precede, and thus trigger, the onset of the transition from the intermediate stage to the close pairing stage (Extended Data Fig. 6e and f).

### RHJ requires Spo11/Dmc1-promoted recombination

In budding yeast, as in most organisms, homolog pairing is mediated by recombination (above). Correspondingly, RHJ is completely abrogated by a catalysis-defective mutation of Spo11 (*spo11y135f*), the transesterase that creates the DSBs that initiate recombination (no events among 11 cells; e.g. Fig. 3a). All subsequent events, including initial nascent D-loop interaction with a partner duplex, ensuing crossover/noncrossover differentiation, and crossover progression, are mediated by the RecA homolog Dmc1 and its meiosis-specific cofactors, including the Hop2/Mnd1 complex ^15^. Correspondingly, absence of either Dmc1 or Hop2 completely abolishes RHJ in ∼80-85% of cells (Fig. 3b and c, n=27 and 14 cells respectively). Interestingly, however, in both mutants, in ∼15-20% of cells, pairing does progress to the intermediate stage, but close pairing does not occur (Fig. 3d, average distance 0.82 µm for *dmc1Δ* and average 0.79 µm for wild type, n=5 and 14 cells respectively, Kolmogorov-Smirnov test *P* = 0.12). In these cases, the intermediate stage is likely achieved by residual strand exchange activity of mitotic RecA homolog Rad51 ^16^ while, in wild type meiosis, Dmc1 is specifically required for later steps.

**Fig. 3.**
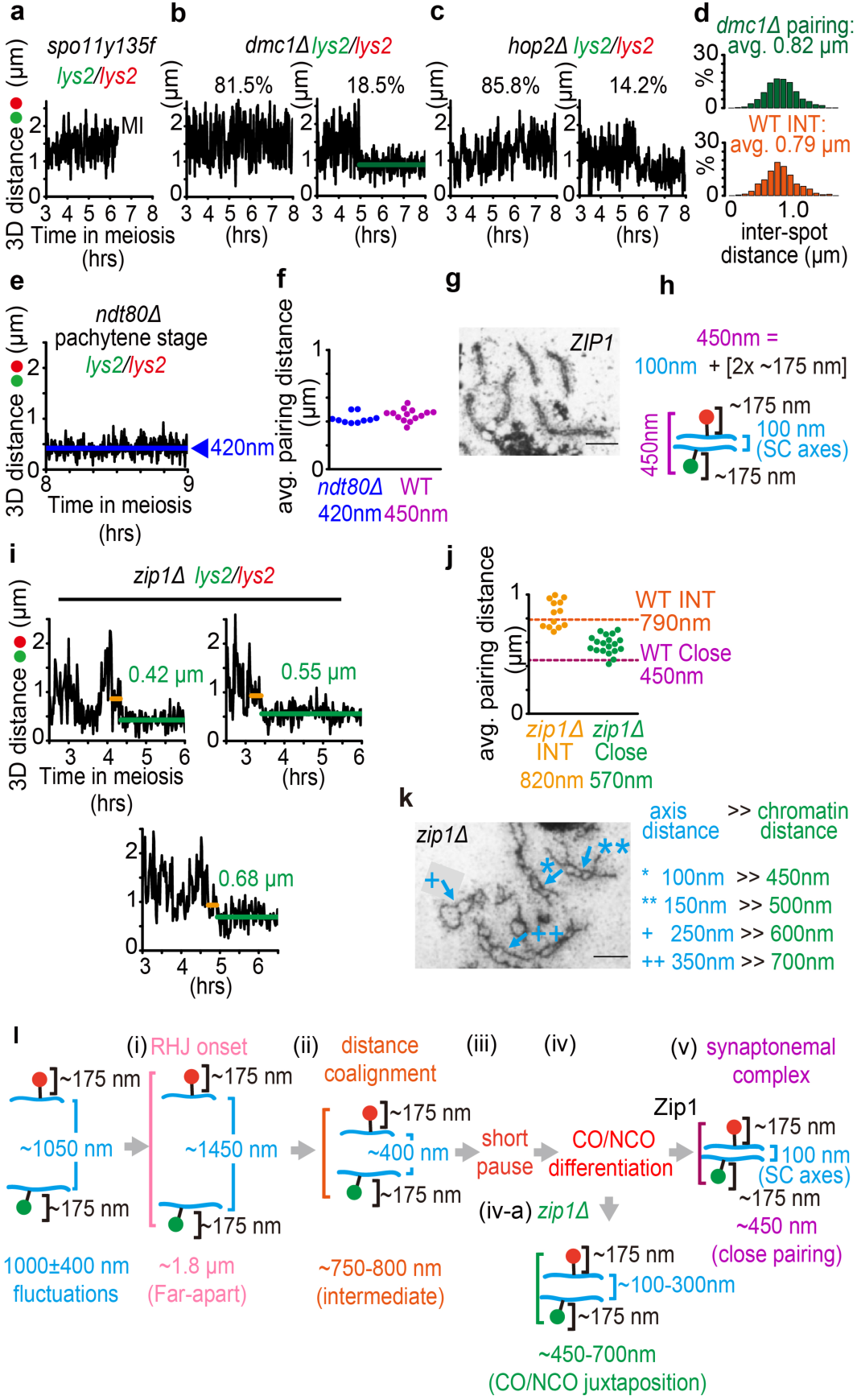
Relationship of RHJ stages to canonical prophase pathway. **a-d.** RHJ requires Dmc1-promoted strand invasion. **a.** Pairing is never detected in *spo11y135f* (n=11 cells). **b, c.** In *dmc1Δ* (**b**) or *hop2Δ* (**c**), most cells show no pairing (left panels), but some cells show partial pairing (right panels) (n=27 cells and n=14 cells total, respectively). **d.** Pairing in *dmc1Δ* progresses to the intermediate stage seen in wild type (green, top versus orange, bottom, n=5 and 14 cells, respectively). **e-h.** Close pairing during RHJ corresponds to the pachytene SC stage. **e, f.** Inter-locus distances in pachytene-arrested cells (**e,** *ndt80Δ*; imaging at 10-second intervals; blue line is average) are the same as for wild type close pairing (**f**; n=10 and n=14 cells, respectively). Thus, close pairing corresponds to SC nucleation/synapsis. **g.** In pachytene SCs, axes are separated by ∼100 nm ^4^. Scale bar = 1 µm. **h.** Thus, ∼450nm spot separation at this stage implies that tagged homolog loci at this stage are located ∼175nm to either side of the SC. **i, j.** In *zip1Δ*, locus pairing progresses to the intermediate stage (orange) and then beyond (green) but with variable final inter-locus distances that are usually greater than for wild type close pairing. Examples in (**i**) and summary statistics in (**j**) at *lys2* (n=19 for *zip1Δ* cells). **k.** (left) In *zip1Δ*, homolog axes are coaligned but with greater separation than in wild type SCs (Extended Data Fig. 7) ^4^. (right) Axis separation distances in *zip1Δ* predict the observed spot separation distances (green) for tagged loci to compare with panel **j**, right. Scale bar = 1 µm. **l.** Steps of RHJ.

### In wild type meiosis, the intermediate stage corresponds to distance coalignment while close pairing corresponds to the presence of SC

The close pairing stage, once established, is maintained until the MI division (above). This feature suggests that it corresponds to formation of synaptonemal complex, which is present at mid/late prophase and disappears prior to MI segregation. This inference is confirmed by analysis of an *ndt80Δ* mutant in which meiosis is arrested at the pachytene stage with full length SC: inter-spot distances are identical in the mutant and for close pairing in wild type (average 0.42 µm vs 0.45 µm; Kolmogorov-Smirnov test *P* = 0.06, Fig. 3e and f, n=10 and 14 cells, respectively, Supplementary Video 4).

Within the SC, homolog axes are linked by central region components at a distance of ∼100 nm with homolog chromatin positioned to either side (Fig. 3g) ^4^. We can therefore infer that, at this stage, the fluorescent tags analyzed in the present study are located, on average, ∼175 nm from their respective axes (Fig. 3h; [(∼450 - 100) = (350)]/2 = 175)]. Based on these findings, in other situations where chromatin is expected to be external to their homolog axes, (average) inter-spot distances can be inferred to be ∼350nm more than inter-axis distances. Most notably, the intermediate stage, characterized by inter-spot distances of ∼800 nm (above), should correspond to a state where axes are separated by ∼400 nm (Fig. 3l (ii), below). This inter-axis distance corresponds directly to that which defines the canonical distance coalignment stage, which is manifested in budding yeast as pairs of Dmc1/Rad51 foci separated by the appropriate distance ^17^.

### Transit from the intermediate stage to close pairing is mediated by crossover/noncrossover differentiation (crossover interference)

In budding yeast, SC formation is nucleated at recombination sites that have undergone crossover/noncrossover differentiation, primarily at crossover-designated sites ^3,5–7^. Correspondingly, since RHJ includes SC formation (above), it should also include crossover/noncrossover differentiation. And in *Sordaria macrospora*, this process comes into play at the intermediate/distance coalignment stage (Fig. 1a) ^18,19^. Together these considerations suggest that, during RHJ, crossover/noncrossover differentiation acts at the intermediate (coalignment) state and triggers progression to close pairing (synapsis).

This inference is supported by analysis of RHJ in a mutant that lacks the main SC central region transverse filament protein Zip1 but still exhibits crossover interference ^6^. We find that, in this mutant, RHJ progresses normally to the intermediate stage and then, after the typical short delay, progresses to a closer distance (e.g. Fig. 3i and j). However, this closer juxtaposition rarely reaches the normal “close pairing” distance (∼450 nm); instead, it reaches an average of ∼570 nm, with considerable fluctuation around that value (450-700 nm; Fig. 3i and j). Moreover, this phenotype corresponds exactly to the *zip1Δ* phenotype defined cytologically ^4^. By the logic above, the range of inter-spot distances of 450-700 nm seen for RHJ corresponds to a range of inter-axis distances of 100-350 nm. This is exactly the range of inter-axis distances observed in *zip1Δ* by electron microscopy (EM) analysis (Fig. 3k; Extended Data Fig. 7) ^4^. These findings imply that: (i) crossover/noncrossover differentiation indeed acts at the intermediate stage; (ii) is the rate-limiting step for exit from that stage; and (iii) results in closer axis juxtaposition. However, (iv) Zip1 is required to achieve the normal final close pairing distance.

Two details of the *zip1Δ* phenotype remain to be defined. (i) The closer juxtaposition observed during RHJ and by EM could reflect effects exerted only at crossover-designated sites, with effects on adjacent regions due to axis stiffness, or could include events at both crossover- and noncrossover-designated sites. (ii) Zip1 could be required for full close RHJ because of its role in SC nucleation; however, it is also possible that Zip1 plays a role in juxtaposition which precedes and is then required for SC nucleation *per se*, as seen in *Sordaria macrospora* ^19^.

### Components of RHJ

Taken together, the above findings define the sequential stages of RHJ (Fig. 3l): (i) Onset initiates by motion-mediated extension of a nascent inter-homolog linkage to an inter-spot distance of 1.8 µm, corresponding to a predicted inter-axis distance of ∼1.4 µm. (ii) Juxtaposition progresses rapidly to an intermediate stage corresponding to canonical ∼400 nm distance axis coalignment. (iii) Pairing pauses for several minutes. (iv) Crossover/noncrossover differentiation then occurs, giving further juxtaposition (as seen in *zip1Δ* (iv-a) and, in wild type meiosis, (v) a concomitant final juxtaposition mediated by Zip1/SC.

### RHJ occurs after prophase structure has developed

Meiotic chromosomes develop prophase organization of cooriented sister linear loop arrays (Fig. 1b). To define the timing of this event relative to the timing of RHJ, we defined the dynamics of chromosome shortening and homolog pairing simultaneously using fluorescent tags at pairs of well-separated loci (e.g. Fig. 4a and b, top, Extended Data Fig. 8a). Development of longitudinal organization is reported by the distance between adjacent loci along a given homolog. This distance undergoes a discrete decrease at ∼2.5 hours after initiation of meiosis. Once achieved, this chromosome length remains constant until onset of Meiosis I, i.e. through pachytene (e.g. Fig. 4a and b, middle; 4c). Thus, the observed shortening transition corresponds to the development of the mature mid/late prophase chromosome organization (Extended Data Fig. 8b). Occurrence of pairing is reported by the distance between tagged loci at corresponding positions on homologs. RHJ onset always occurs well after the shortening transition (e.g. Fig. 4a and b bottom versus middle and 4c). The average delay observed was ∼28 min (Fig. 4d, 4 types of strains, 31 cells).

**Fig. 4.**
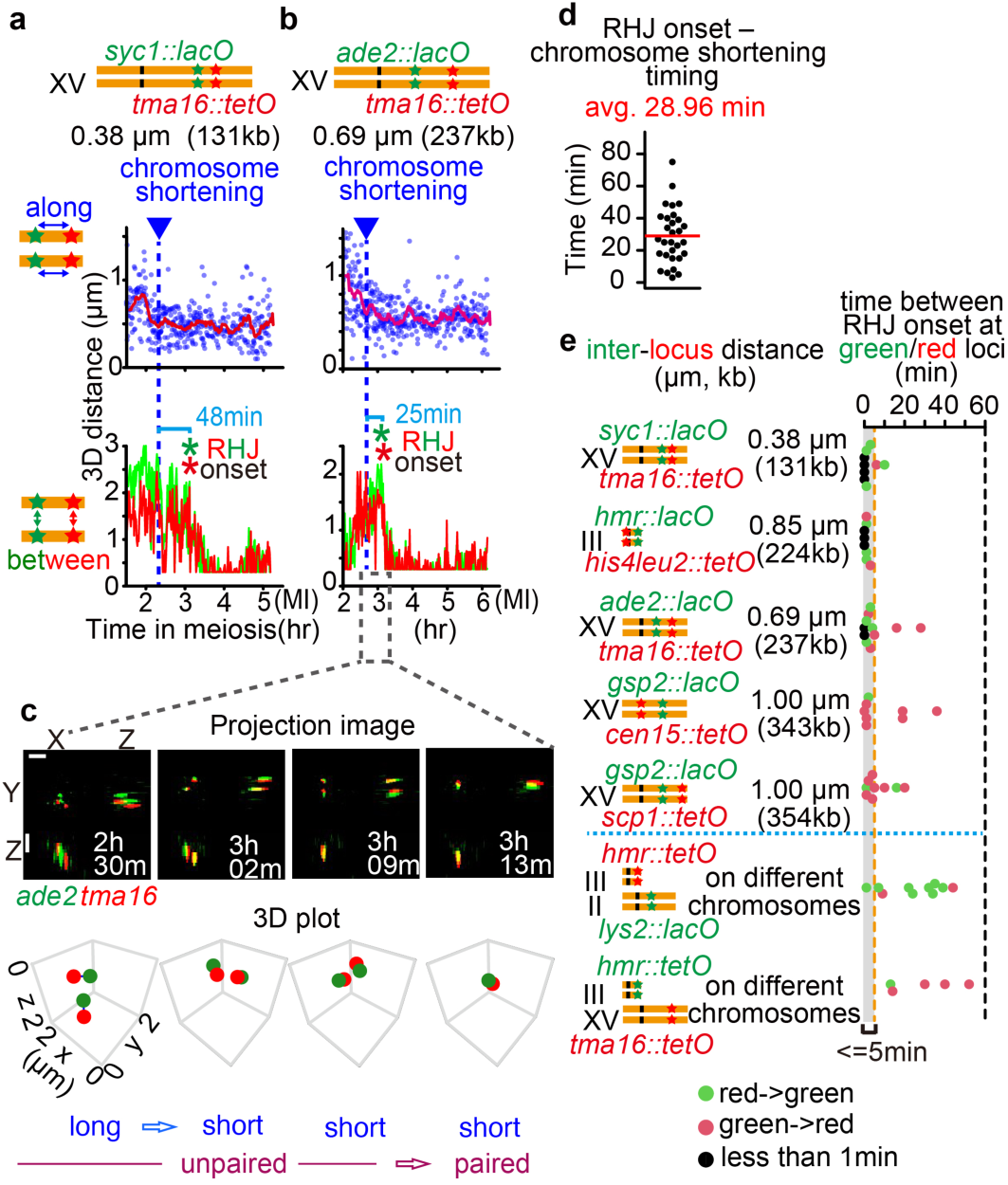
RHJ at adjacent loci is temporally coupled along well-organized homolog chromosomes. **a, b.** Top: two versions of Chromosome XV carried the indicated labels at pairs of loci separated by the indicated distances which represent ∼15-25% total chromosome length, respectively. Physical distances are based on pachytene chromosome lengths ^36^. Middle: variations in red/green distance report longitudinal compaction. Red line is the average distance in a 10 min sliding window. Vertical blue dotted line indicates the time when chromosomes shorten to a state that persists into pachytene. Bottom: green/green and red/red spot distances report homolog locus pairing. Onset (*), defined as in Fig. 2h, occurs well after shortening. Resolution of same color spot distances is limited to 300 nm. **c.** Projected images (top) and 3D plots (bottom) for selected images from the indicated time period of panel **b**. Scale bars = 2 µm. **d.** Distribution of time differences between RHJ onset and chromosome shortening at *syc1-tma16*, *ade2-tma16*, *gsp2-cen15*, and *gsp2-scp1* (n=31 cells; red bar indicates average). **e.** Five pairs of adjacent loci (top) and two pairs of loci on different chromosomes (bottom) were examined for the time interval between green/green and red/red locus pairing as in (**a, b**). Dashed orange and black vertical lines define an interval of 5 min and 60 min, respectively. Different colored dots indicate which locus pairs first (green or red) or if the interval is less than one minute (black).

### RHJ occurs nearly simultaneously at adjacent loci separated by 300kb (∼1 µm), but occurs asynchronously throughout the genome

In both of the examples shown in Fig. 4a, b, the two adjacent loci undergo RHJ nearly simultaneously. This effect is general, as shown by systematic analysis of pairs of adjacent loci located at varying distances and positions along short and long chromosomes (Fig. 4e, Extended Data Fig. 8c-f, Supplementary Video 5 and 6). In 35/45 such comparisons, the two sets of homolog spots undergo RHJ onset within five minutes of one another, although differences of up to ∼40 min were sometimes observed (Fig. 4e top). The pairs analyzed in these cases were separated by distances of up to ∼350 kb, corresponding to ∼1 µm chromosome length, which corresponds to about a third (or more) the length of an average budding yeast chromosome. Loci separated by greater distances have not yet been examined. In contrast, RHJ onset at homolog loci on two different chromosomes is usually separated in time by more than 10 min (12/15 cases) and occurred within five minutes in only 1/15 cases (Fig. 4e bottom, Extended Data Fig. 8g and h).

Interestingly, two loci on unrelated chromosomes can undergo RHJ at times differing by up to ∼50 min (Fig. 4e bottom, Supplementary Video 7). This difference implies that RHJ occurs asynchronously in different regions throughout the genome, with a total “pairing period” of about an hour required for RHJ at all loci. This pattern is consonant with the known timing of events in yeast meiosis (∼1-2 hours from DSB formation to SC formation) ^20^ and the nearly universal finding, from “snapshots” of meiotic nuclei in diverse organisms, that, at any given time, different regions of the genome are at different stages of the pairing program. Progression through the genome may be partially hard-wired: in budding yeast and mice, the timing of recombination initiation is directly correlated with the timing of DNA replication on a region-by region basis ^21,22^.

### End-led motion promotes both onset and progression of RHJ

Cytoskeleton-mediated telomere-led movement is prominent in budding yeast. This motion requires meiosis-specific protein Ndj1, which links telomeres to the nuclear envelope-spanning “LINC” complex, one component of which is Csm4. The LINC complex in turn is connected to the motor protein Myo2, which moves the entire ensemble along actin filaments ^8,14^.

Analysis of a tagged *lys2* locus shows that motion initiates during meiosis, prior to RHJ, and continues into pachytene, thus after completion of RHJ. This is seen by the total space explored in 5 min (Fig. 5a left) and the distance traveled in 10 sec and 1 min (Fig. 5a right, Extended Data Fig. 1d and g). Absence of Ndj1 or Csm4, or treatment with actin depolymerizing agent Latrunculin B (LatB) progressively decreases motion, as seen by both criteria (Fig. 5a). These treatments also reduce the extent of fluctuation of homolog inter-spot distances at *lys2* as defined by the Mean Square Change in Distance (MSCD) (Fig. 5b).

**Fig. 5.**
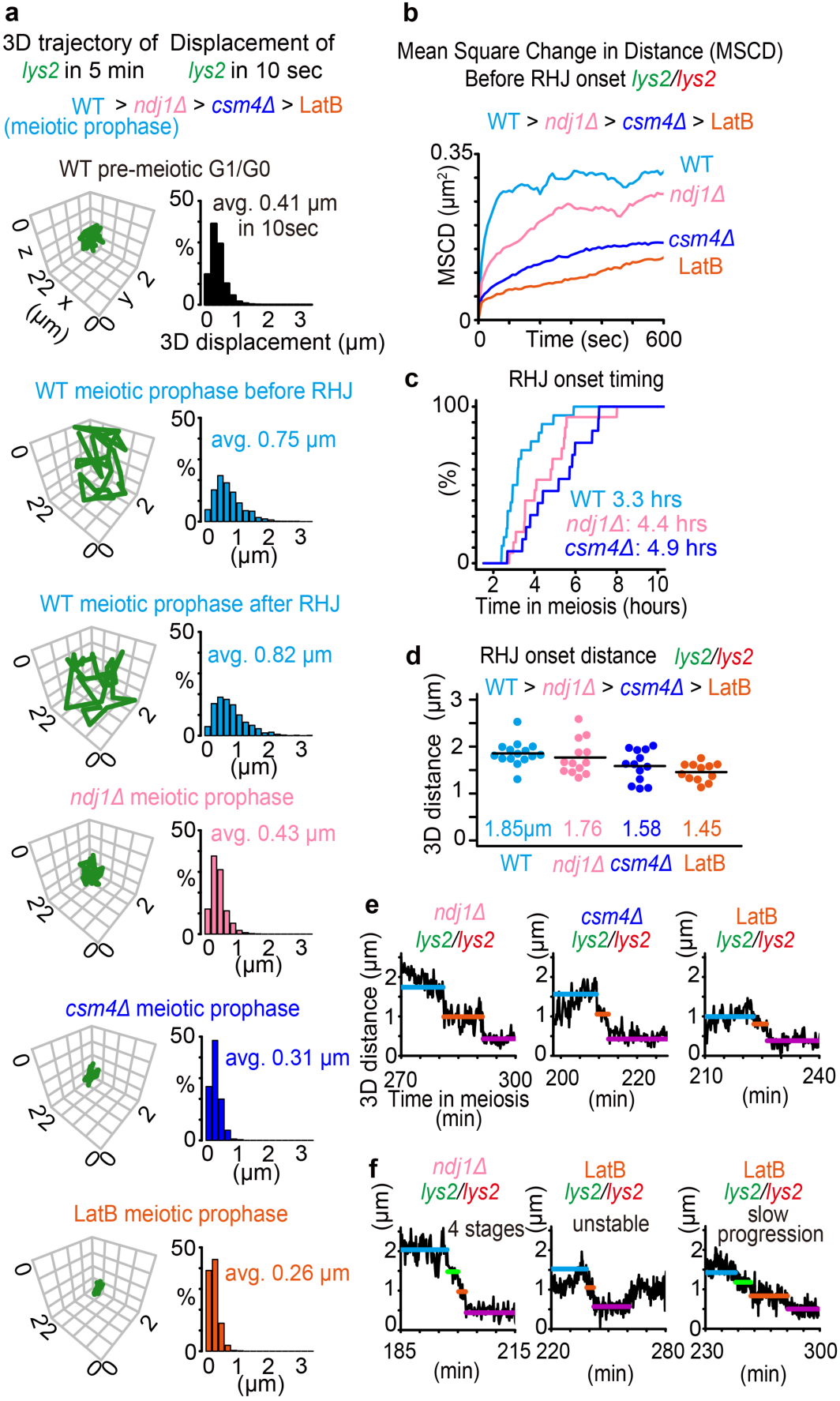
Disruption of rapid chromosome motion affects RHJ onset. **a.** Motion of individual *lys2* spots in the indicated conditions as defined by a representative 5 min trajectory (left) and the average displacement in 10 sec from many individual cells (n= 10-15 cells). **b-d** Correlated effects of different perturbations on inter-spot motion and the timing and distance of RHJ onset, defined at *lys2* (n=10-18 cells). **b.** Mean square change in distance (MSCD) between homologous *lys2* spots during the 20 mins prior to RHJ onset. **c.** Time of RHJ onset after transfer of cells to meiotic medium, defined at 1 min resolution. **d.** Distribution of the distances at the RHJ onset in individual cells (black line = average). **e, f.** Cells in which motion has been perturbed sometimes show normal progression of RHJ (**e**) and sometimes show abnormal progression (**f**, see the details in Extended Data Fig. 9b). Far-apart state (before RHJ, cyan); intermediate state (orange) and close pairing (purple) as in wild type (Fig. 2a, b); abnormal progression sometimes shows an extra state (green).

The hierarchy of effects is observed by all criteria. Motion in *ndj1Δ* resembles that in mitotic cells, in accord with complete abrogation of the entire meiotic apparatus due to absence of meiosis-specific telomere/nuclear envelope association (Fig. 5a). *csm4Δ* exhibits less motion than *ndj1Δ*; and motion in LatB is most severely reduced (Fig. 5a, b). In these latter cases, telomere/nuclear envelope associations are still intact but are, apparently, less mobile in *csm4Δ*. In the presence of LatB, chromosomes and their ends are virtually motionless (see also ref 12). These phenotypes have the interesting implication that meiotic telomere/nuclear associations are a double-edged sword: on the one hand, they likely help to avoid entanglements by keeping ends from threading among different chromosomes; on the other hand, they render the interhomolog interaction program strongly dependent on cytoskeleton-mediated end-led motion.

All three motion-defective conditions impede RHJ onset, in two respects.

- First, the time of pairing onset relative to initiation of meiosis as defined by long timescale imaging, is delayed in *ndj1Δ* and *csm4Δ* (Fig. 5c). Importantly, this delay arises after initiation of recombination because *ndj1Δ* and *csm4Δ* cells exhibit normal DSB timing ^23^. Analogously, in LatB-treated cells, long timescale imaging is problematic, but relevant time points exhibit a highly elevated fraction of cells with unpaired spots, implying a severe defect (Extended Data Fig. 9a).
- Second, all three conditions reduce the characteristic RHJ “onset distance” (Fig. 5d). The severity of both effects is again correlated with the severity of motion reduction (wild type < *ndj1Δ* < *csm4Δ* < LatB; Fig. 5c, d). These findings are concordant with the fact that, in wild type meiosis, motion directly triggers RHJ onset (Discussion).

In all three motion-defective situations, RHJ *per se* still occurs and often proceeds with normal kinetics (Fig. 5e, Supplementary Video 8). However, a significant proportion of nuclei (17/38 in the perturbed conditions versus 1/15 in wild type for *lys2* loci, Fisher’s exact test *P*=0.009) exhibit atypical progression, e.g. discrete pauses with an extra step, unstable pairing or slower progression of RHJ (Fig. 5f; Extended Data Fig. 9b). Perhaps chromosome “jostling” is helpful in moving unrelated chromosomes out of the way, thus removing impediments to RHJ progression.

## Discussion

Analysis of 3D dynamics during homolog pairing, in real time, at intervals of 10 sec, has revealed the existence of “rapid homolog juxtaposition” (RHJ). Corresponding loci on homologs come together in space by a multi-step sequence of events (Figure 3l) which usually occurs in the span of just a few minutes. This progression is initiated when homolog loci are separated by a considerable distance. The two loci first come rapidly together into a discrete intermediate configuration corresponding to distance coalignment. Then, after a brief pause, crossover/noncrossover differentiation (interference) promotes further juxtaposition, which is concomitantly finalized by effects of Zip1/SC nucleation.

We further find that: (i) RHJ occurs nearly synchronously over substantial chromosome lengths, while occurring asynchronously throughout the genome; and (ii) cytoskeleton-mediated end-led motion is important for both onset and progression of the RHJ program.

### Motion-mediated RHJ onset

The recombination interaction present at the time of RHJ onset presumably comprises a nascent D-loop linkage (Fig. 1b). However, the detailed molecular state of this linkage is not known. Since chromosome axes have developed at the time of RHJ onset, the nascent D-loop complex might already be associated with the partner axis at the time of onset, although this remains to be established. The DNA component of the linkage could be a single RecA homolog-coated DSB end on a long double-strand DNA “tentacle”, an axis-associated RecA/ssDNA filament associated with a partner chromatin loop sequences and/or some other structure ^20,24,25^.

RHJ is triggered by a motion-mediated increase in inter-spot distance. An interesting possibility is that this increase creates mechanical tension along the nascent inter-homolog linkage. For example, tension might trigger an axis-associated recombination complex/axis ensemble to initiate the juxtaposition process. It is also interesting that reductions in end-led motion delay both RHJ onset and the distance at which onset occurs, both of which are altered in correlation with the extent of motion reduction. Perhaps onset can eventually be triggered at a shorter distance in these cases, but with a reduced probability per time.

### Rapid, directed juxtaposition into the coalignment state

Identification of a discrete intermediate pairing state corresponding to distance axis coalignment matches the cytological observation of such a state in diverse organisms (Introduction) and confirms previous evidence for occurrence of this state as defined by separated pairs of Rad51/Dmc1 foci in budding yeast ^17^. The time required for progression from RHJ onset to this intermediate stage is very short (∼30 sec). During this transition, homolog loci approach one another in a smooth, continuous fashion, at a rate faster than that seen for earlier motion-mediated fluctuations (Extended Data Fig. 5). These features imply operation of an active process, the nature of which remains to be established. Possibilities include loop formation/extrusion, condensate formation or motor protein translocation, all of which can occur at appropriate rates ^12,26–29^, or some unknown process.

### Crossover/noncrossover differentiation triggers juxtaposition to close pairing

The finding that crossover/noncrossover differentiation (as governed by crossover interference) mediates the second RHJ transition is interesting from several perspectives. First, it reveals that crossover patterning decisions act at the intermediate/coalignment state in budding yeast, as in Sordaria (Introduction). Second, it reveals that this process has the structural effect of bringing homolog axes closer together, independent of SC nucleation. We do not yet know whether this further juxtaposition occurs only at sites of crossover differentiation or at additional (noncrossover) sites. Third, this second transition of RHJ is similar in nature to the first transition, into the intermediate state: it is also smooth and continuous, occurs at a similar rate, and might also be triggered by an increase in inter-spot distance. Perhaps the two transitions are mechanistically related. In Sordaria, this transition involves formation and contraction of DNA/recombination/structure bridges ^19^, by unknown mechanisms. Fourth, since the duration of the intermediate state essentially defines the overall duration of RHJ, occurrence of crossover/noncrossover differentiation is the rate-limiting step for completion of homolog pairing. Fifth, since the duration of the intermediate stage is usually just a few minutes, crossover/noncrossover differentiation follows close on the heels of coalignment. Finally, since exit from the intermediate stage defines the timing of crossover/noncrossover differentiation, this event provides a diagnostic assay for a very early step of the differentiation process, thus allowing future analysis of the timing of that event, across the genome, in wild type and mutant situations.

### Implications for global pairing

The present study examines homolog juxtaposition at individual tagged loci and defines the corresponding local steps. However, several of the features defined for RHJ could potentially have important global effects, i.e. in helping to minimize formation of entanglements among unrelated chromosomes and inappropriate (ectopic) recombination between non-homolog chromosomes ^1,30^.

- RHJ onset requires that corresponding homolog regions are at a distance allowing motion-mediated fluctuations to give the requisite extension. One aspect of this requirement is that homologs cannot be too far apart. RHJ onset occurs at a distance of 1.8 µm (tagged loci) or ∼1.4 µm (axes). In yeast, this is about 60-70% of the diameter of the nucleus (∼2.5-3 µm), and end-led motion involves appropriate displacements ^12^. However, there are hints of long-range interactions of similar length in plants and mice ^25,31,32^, and cytoskeleton-mediated motions involve similar mechanisms in all organisms and thus might be expected to be similar in magnitude in all cases. If the same RHJ onset requirement pertains generally across organisms, it would be a powerful mechanism for ensuring that homologs were already in a joint space before the dramatic events of juxtaposition are initiated. Tight linkage of RHJ onset to motion also fits with the suggestion that back-and-forth motions of different groups of telomeres could be a critical factor in placing homolog regions in joint spaces ^1,33^.
- Sensing of extension to trigger RHJ onset also requires an unimpeded connection between homologs. The requirement for extension thereby ensures close, stable pairing will not be initiated if an unrelated chromosome is in the way.
- The fact that RHJ onset occurs nearly simultaneously at loci separated by thirds (or more) of chromosome lengths raises the question of whether onset of juxtaposition at one locus might promote onset at adjacent loci. Such an effect could occur simply by virtue of axis stiffness ^34^. An alternative/additional possibility is raised by the fact that early recombination (“crossover precursor”) interactions are evenly spaced, as seen and/or inferred for both budding yeast and Sordaria ^34–36^ and for certain early recombination interactions in mouse ^37^. This effect cannot be achieved by independent initiation of interhomolog interactions on the two homologs and thus points to the existence of a patterning process that occurs prior to imposition of classical crossover interference (e.g. ref 37). Perhaps RHJ onset is responsible for such patterning. For example, an RHJ event at one locus might simultaneously encourage occurrence of another event nearby while, at the same time, ensuring that the nearby event occurs at some distance away. This effect could potentially act at the DSB level and thereby underlie the “DSB interference” seen along individual chromatids ^38^.
- Crossover interference chases RHJ-mediated coalignment through the genome. This coupling could be important for minimizing entanglements among unrelated chromosomes. Such entanglements are trapped largely by recombination-mediated coalignment interactions ^1^. And crossover interference has the effect of eliminating interhomolog connections at sites that are designated to be matured as noncrossovers. Since crossover/noncrossover differentiation follows closely on the heels of coalignment, on a locus-by-locus basis and coordinately along the chromosomes, potentially deleterious constraining interactions will be eliminated dynamically in concert with coalignment and juxtaposition progresses throughout the genome.

## Summary

Corresponding loci on homologs undergo motion-triggered rapid juxtaposition to and through coalignment, crossover interference and SC formation, all in the space of a few minutes, with interesting implications for both local and global events.

## Acknowledgements

We thank members of the Kleckner laboratory and D. Zickler for advice and discussions. This research, T.N., B.W., and N.K. were supported by a grant to N.K. from the National Institutes of Health (R35 GM136322). T.N. was supported by fellowships from Japan Society for the Promotion of Science and Charles A. King Trust.

## Author contributions

T.N. and N.K. conceived and designed experiment. T.N. and B.W. constructed the strains. T.N. performed imaging and data analysis. T.N. and N.K. wrote the article. All authors contributed to the article and approved the submitted version.

## Materials and methods

### Yeast strains

*Saccharomyces cerevisiae* strains are isogenic derivatives of wild-type SK1. *TetO* array (∼120-240 repeats)/TetR-tandem-mCherry and *LacO* array (∼120-240 repeats)/LacI-mEGFP (A206K) systems are based on Lau et al. 2003 (ref 39). Tandem-mCherry was created based on the mCherry sequence with the linker sequence used in tdTomato ^40^. Genotypes of strains are given in Supplementary Table S1.

### Meiotic time course

All operations were performed at 30°C. Strains maintained in glycerol stock at –80°C were patched onto YEPG plates (3% w/v glycerol, 2% w/v bactopeptone, 1% w/v yeast extract, 2% w/v bactoagar) overnight at 30°C. Cells were streaked out to single colonies on YEPD plates (2% w/v bactopeptone, 1% w/v yeast extract, 2% w/v glucose, 2% w/v bactoagar) and grown for two days at 30°C. A single colony is transferred to 4 ml YEPD liquid medium (2% w/v bactopeptone, 1% w/v yeast extract, 2% w/v glucose) and grown overnight at 30°C. A 1/100 dilution of the culture was made with YEPA medium (1% w/v potassium acetate, 2% w/v bactopeptone, 1% w/v yeast extract, 2 drops per liter antifoam) and grown for 13.5 hours in a 2L flask with vigorous shaking at 30°C. Meiosis was induced by the transfer of cells to 0.3 % SPM (0.3 % w/v potassium acetate, 0.02% w/v raffinose, 2 drops per liter antifoam) in a 2 L flask with vigorous shaking at 30°C. Upon transfer to SPM, the cells initiate meiosis synchronously and efficiently ^20^. To induce the CUP1 promoter, 50 µM of CuSO_4_ was added to SPM at the timing of the media change.

### Chemical treatment

Latrunculin B (LatB, Santa Cruz Biotechnology) was dissolved in dimethyl-sulfoxide (DMSO). LatB was added to a final concentration of 30 μM at 2.5 hours after transferring cells to SPM, and imaging started at 3 hours in SPM (t = 3 h of meiosis).

### Live-cell imaging

Cell samples from an experimental culture were vortexed and 2 µl of cell samples were quickly spread onto a glass base dish (Matsunami) coated with Concanavalin A. A premade agarose pad (1%, soaked in SPM media for a day) was placed gently over the liquid drop of cells, and excess media was absorbed by a piece of Kimwipe. Cells were observed at 30°C kept by the incubator system (In Vivo Scientific) using a Ti microscope (Nikon) equipped with 488/561 dual-color filters (Semrock, Brightline, LF 488/561 - 2X2M - BNTE) for the Pinkel configuration, an sCMOS camera ORCA-Flash 4.0 (Hamamatsu Photonics), and a piezo device (Physik Instrumente) for acquiring Z stacks. Cells were exposed to the LED light (Lumencor) through an objective lens (60 × PlanApo, NA 1.40; Nikon). The microscopy system was controlled, and images were acquired, through µ-Manager software and MATLAB (MATLAB 2014b) ^41^. For live-cell imaging, movies of 13 z-stack steps with 389-nm step size were acquired using µ-Manager software with 10 ms exposure time and 16.6 ms camera acquisition time. Z-stacks were acquired at 1-minute or 10-second intervals for 10 hours or 2 hours, giving a total of 601 or 721 Z-stack images. After taking Z-stack images for fluorescence, the brightfield images with 13 Z-stack were obtained to determine the cell position. Imaging was started after 1.5 hours incubation in SPM media for long-timescale imaging and after 2.5 hours incubation in SPM for short-timescale imaging. For short-timescale imaging of *ndj1Δ* and *csm4Δ* mutant cells and LatB-treated cells, imaging was started after 3 hours of incubation in SPM because of the delay of pairing timing. For short-timescale imaging of G1/G0 cells, imaging started just after transferring the cells from YPA to SPM. For short-timescale imaging of *ndt80Δ* mutant cells, imaging started after 8 hours of incubation in SPM to observe the cells at the pachytene stage.

### Data analysis

#### Low-SNR data processing

Prior to analysis, images were denoised using a custom MATLAB program (A1 filtering) that accurately defines the presence and positions of fluorescent spots at low signal-to-noise ratios ^12,42^. This algorithm dramatically extends the ability to capture many images over long periods of time, as illustrated in this work (Extended Data Fig. 1).

#### Spot detection and tracking

After A1 filtering and cell segmentation by the above program (Extended Data Fig. 1a), the center positions of spots were detected and tracked by ImageJ Fiji plug-in TrackMate ^43^. After spot detection by TrackMate, every spot was manually checked, and corrected when necessary (∼1% of total spots). In general, when each homolog was labeled with a single operator array (e.g. Fig. 1c and e), each nucleus contained only a single fluorescent focus of each color. As such, we encountered no problems with spot tracking. 3D projection and kymograph plots (e.g., Fig. 1e, f) were generated by a homemade program ^42^, and 3D plotting of the spot and trajectory was performed by plot3D package in R (e.g., Fig. 1g bottom). The 3D projection movie was created based on the image of 3D projection after applying “Brightness/Contrast Adjustment” in Fiji (e.g., Supplementary Movie 1). For analysis of long-timescale imaging of wild type cells (e.g. Extended Data Fig. 1b-g), only cells that went through meiosis normally (premeiotic G1/G0 stage at 1.5 hours in SPM, rapid chromosome motion, and MI division in ∼10 hours) were analyzed. For analysis of long-timescale imaging of mutant cells (*spo11yf*, *dmc1Δ*, *hop2Δ*, *zip1Δ, ndj1Δ,* and *csm4Δ*; Fig. 3a-d, 3i, 3j, 5c), the only cells that seemed healthy (MI, rapid chromosome motion, or pairing was observed) were used. For the analysis of the short-timescale imaging, the cells that exhibited pairing during the imaging were used and analyzed. The pairing ratio of LatB-treated cells was determined by the kymograph of the cells. Spot detection accuracy in this study was based on our previous work using the same imaging and analysis system (∼40 nm in the X/Y dimensions and z: ∼75 nm in the Z dimension) ^12^. In this previous study, the spot detection accuracy was estimated by the fluctuation of spot detection in continuous imaging of the fluorescent spots in the cells fixed by paraformaldehyde. In this study, two-colored imaging further allows the assessment of inter-spot distance not constrained by the diffraction limit.

#### Calculation of 3D distances between detected fluorescent spots

##### Single green and single red spots

The 3D distance between the green spot (*x_g_, y_g_, z_g_*) and the red spot (*x_r_, y_r_, z_r_*) was calculated by ((*x_g_* – *x_r_*)^2^ + (*y_g_* – *y_r_*)^2^ + (*z_g_* – *z_r_*)^2^) ^½^. When the green spot or red spot was rarely and temporarily split into two spots, the midpoint of the two spots was used as the spot position. *Multiple spots (2 green spots and 2 red spots)*: In the case of the multi-spot labeling (e.g., Fig. 4a, b), two categories of distance were calculated. First, for the calculation of the distance between spots along the chromosomes (green vs. red), two combinations could be chosen when we could observe all four spots (Extended Data Fig. 8a). The combinations including the minimum of maximum distances were estimated as the spot distances along each chromosome (Fig. 4a, b, middle, Extended Data Fig. 8a). When we could see only one spot vs. two spots or one spot vs. one spot for the green vs. red spot(s), we calculated the distance(s) for every combination(s) of the spot(s). Second, to calculate the distances between fluorescent spots of homologous loci (green vs. green and red vs. red), the calculation of the distance was the same as with single green/red spots analysis described above (e.g., Fig. 4a, b bottom). When only a single spot was detected, we set 300 nm as the distance because of the diffraction limit. When the distance was below 300 nm, 300 nm was also set as the distance.

#### Distance change analysis

Distance change analysis is summarized in Extended Data Fig. 3.

##### Stages and steps detection

After calculating the distances, the cpt.mean function in the changepoint package of R ^44^ was used to detect the stages of homolog pairing (Fig. 2a). The 3D distance plot was divided into three stages. Three stages were defined as far-apart (before RHJ), intermediate, and close pairing (Extended Data Fig. 3a right). These transitions were also very prominent, and these transitions were estimated by the number of change points = 2 or 3 in ∼60 % of the cells. In other cells, the number of change points that could detect the intermediate stage during the transition from far apart to close pairing was searched and applied. In some cases, the stages at the transition from before pairing to pairing could not be estimated as three stages, but as two or four stages (Extended Data Fig. 2c and d). When there was no intermediate stage between the ‘far apart’ and ‘close pairing’ states, we determined the RHJ pattern as 2 stages (single step RHJ). When the obvious four stages were detected and transitions between each stage were smooth, we determined the RHJ pattern as 4 stages (e.g., Fig. 5f left and right, Extended Data Fig. 2d). For the unstable close pairing, when the obvious and temporary recoiling of the distances to the intermediate stage after the close pairing initiation was observed, we determined the RHJ pattern as unstable close pairing (e.g., Extended Data Fig. 2e).

##### Inter-spot distance distribution

After the detection of the start and end points of the intermediate stage, the distribution of inter-spot distances for the intermediate stage (from the start to the endpoint) was calculated. Then, the distribution of inter-spot distances for the far-apart (before RHJ) stage was calculated for the time point from one frame before the intermediate start to the start point of the spot tracking, and the distribution of inter-spot distances for the close pairing stage was calculated for the period from one frame after the end of the intermediate stage to the end of the spot tracking.

##### Detection of local peak and valley points (local maximum and local minimum)

The onset of RHJ (symbolled by *, e.g., Fig. 2a, Extended Data Fig. 3b) was defined as the local peak (local maximum) prior to the start of the intermediate stage. The local peak was defined by the following process.

i. The searching process was started from the beginning of the intermediate stage (e.g., time *t*=s). The searching process went backward and determined the first point *t*=s-n where d*_t_*_=s-n-1_ < d*_t_*_=s-n_. d*_t_*_=s_ means the distance between homologous loci at *t*=s. d*_t_*_=s-n_ was set as the distance at the temporary peak and *t*=s-n was set as the temporary peak time.
ii. d*_t_*_=s-n_ was compared with d*_t_*_=s-n-1_, …, d*_t_*_=s-n-m_, …, d*_t_*_=s-n-a_. (a = 3 when 10 sec intervals and a = 2 when 1 min intervals). If the time point *t*=s-n-m where d*_t_*_=s-n-m_ > d*_t_*_=s-n_ was found, d*_t_*_=s-n-m_ was set as the new temporary peak, and *t*=s-n-m was set as the new temporary peak time. If there were several points of *t*=s-n-m’ where d*_t_*_=s-n-m’_ > d*_t_*_=s-n_, the maximum d*_t_*_=s-n-m’_ was set as the new temporary peak, and *t*=s-n-m’ was set as the new temporary peak time. After this process, the searching process went back to (ii), and the higher temporary peak was searched.

If *t*=s-n-m where d*_t_*_=s-n-m_ > d*_t_*_=s-n_ was not found in d*_t_*_=s-n-1_, …, d*_t_*_=s-n-m_, …, d*_t_*_=s-n-a_, d*_t_*_=s-n_ was determined as the local peak (local maximum), and *t*=s-n was determined as the local peak time, and the searching process of the local peak ended.

To search the local minimum just after the intermediate end as the RHJ endpoint (local valley symbolled by #, Extended Data Fig. 3b), the same but reverse searching process was used (for local minimum search upward from the time point of the intermediate end) (Extended Data Fig. 3b).

##### Calculation of pairing time

Total pairing time was determined by *t*_valley_ - *t*_peak_ .

##### Calculation of duration of intermediate period

The duration at the intermediate stage was determined by *t*_intermediate___end_ -*t*_intermediate_start_ + 1. If there were only two stages (the single step RHJ, Extended Data Fig. 2c), the duration of the intermediate was set as 0.

##### Calculation of durations of transitions

The durations of transitions were determined by *t*_intermediate_start_ - *t*_peak_ and *t*_valley_ - *t*_intermediate_end_.

##### Calculation of time difference between the peak (RHJ onset) and intermediate start

The time between the immediately-preceding local peak (local maximum) and the intermediate start time was determined by *t*_intermediate_start_ - *t*_peak_. As the control, we calculated the time difference between the time point whose distance was in the range of the intermediate stage distance by chance before the RHJ onset (>0.6 µm and <1.2 µm) and the analogous immediately-preceding local peak (nearest local maximum, > threshold value (1.5 µm, 1.6 µm, and 1.7 µm)). The searching process was based on the local maximum search described above. The searching process was started from the time point (e.g., *t*=s) whose distance was at the range (>0.6 µm and <1.2 µm) before the RHJ onset. The searching process went backward and defined the first point *t*=s-n where d*_t_*_=s-n-1_ < d*_t_*_=s-n_ and d*_t_*_=s-n_ > threshold value (1.5, 1.6, or 1.7 µm). After here, the searching process was the same as the maximum searching process described above and the analogous immediately-preceding local peak was determined. The time differences were calculated between all time points before the RHJ onset whose distances were in the range of the intermediate stage distance (>0.6 µm and <1.2 µm) and their analogous immediately-preceding local peak time points. To treat the same conditions, we also calculated the time difference between the analogous immediately-preceding local peak > 1.5, 1.6, or 1.7µm and the intermediate start point (Extended Data Fig. 6c and d).

##### Calculation of max speed of Δdistance

The max speed of Δdistance in the transition from the far-apart stage to the intermediate start was determined by the maximum change in the distances between RHJ onset* and the start of the intermediate stage. The max speed of Δdistance in the transition from the intermediate stage to the close pairing stage was determined by the maximum change in the distances between the end of the intermediate stage and the RHJ endpoint # (Extended Data Fig. 3b).

##### Analysis of chromosome shortening

To determine the chromosome shortening timing, first, the average distance between spots (green vs. red) along the chromosomes for 10 frames (10 minutes) was calculated (the red line in Fig. 4a and b, middle) and a mix of the single/dual sets of distances calculated from the detected spots was used. In the case of single chromosome analysis, a single set of distances was used (Extended Data Fig. 8b). Using the cpt.mean function in the changepoint package in R (described above), the average distance plot was divided into two (or more) stages. The large change of the distance (the first change in almost all cases) was picked up and the local minimum was searched from the change point by the local minimum searching process described above (a = 2 for 1 min intervals). We determined the local minimum point (local valley) of the average distances between spots along chromosomes as the shortening time point, suggesting sometimes this point was the local valley of the distances along chromosomes, the chromosomes and the chromosome shortening initiated earlier than this point. The time difference between the chromosome shortening and the RHJ onset was calculated by the time difference between the time point of RHJ onset of green or red spots that initiated RHJ earlier and the shortening timing. If the timing of the chromosome shortening was not clear (the fluctuation of the distance between two spots along chromosomes was small), we did not use the cells for the chromosome shortening analysis. For example, the chromosome shortening timing of all the *hmr-his4leu2* spots along chromosome IIIs was not clear.

##### Estimation of the physical chromosome length

The physical length between spots along a chromosome was estimated based on the physical length of spread chromosomes at the pachytene stage (3.2 µm and 1.2 µm for ChrXV and ChrIII, respectively) ^36^.

##### Analysis of zip1Δ pairing data

First, the 3D distances between homolog loci were calculated, and pairing steps were determined by the process described above. The average pairing distance in the *zip1Δ* mutant cell was calculated for data points for 1 hour (61 frames) after the initiation of the close pairing stage (but larger than WT, see Fig. 3j).

##### Analysis of dmc1Δ pairing

The 3D distances between homolog loci by 8 hours in the SPM were calculated. The cells showing RHJ by 6 hours in the SPM were picked as “paired” cells. Using the method described above, the pairing steps were detected. In all pairing cases, there were two stages (the stage before pairing and the intermediate-like stage). When the pairing disruption was observed, the distances before the pairing disruption were used for the calculation.

### Motion analysis

The motion of the green spot from *g_1_*(*x_g1_, y_g1_, z_g1_*) at *t_1_* to *g_2_*(*x_g2_, y_g2_, z_g2_*) at *t_2_* were calculated by ((*x_g1_* – *x_g2_*)^2^ + (*y_g1_* – *y_g2_*)^2^ + (*z_g1_* – *z_g2_*)^2^) ^½^. For G1/G0 data (t=0 hours in SPM media), we calculated the motion of the spots within 30 minutes from the start of the imaging. For the motion of the spots before RHJ, we calculated the motion of the spots within 15 min of the beginning of the imaging (t=2.5 hours) if the pairing did not initiate. If the pairing started within 15 min of the beginning of the imaging, we used the motions of the spots before RHJ onset. For the motion of the spots after RHJ, we used motions of spots in 15 min after the RHJ endpoint. For the mutant cells and LatB-treated cells, we calculated the motions of spots in the first 15 min regardless of the initiation of the RHJ.

### Distance change analysis

The distance change in 10 seconds (1 frame) was calculated by *Δd* = |(*d_i+1_* – *d_i_*)|. *d_i_* is the 3D distance between homologous loci at *t=i*.

Mean square change in distance (MSCD) ^45^ was calculated based on the distance between homolog loci (*d_1_*,…, *d_i_*,…, *d_n_*; *Δt (t_n_ – t_n-1_)* = imaging time interval) by 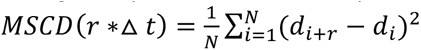.

To calculate the distance change in time for the single trajectory for the RHJ onset to the intermediate start or from the intermediate end to the RHJ endpoint without averaging, a square change in distance was calculated by (*Δ*_012_ − *Δ*_0_)^4^ (Extended Data Fig. 5b).

### Electron microscopy image analysis

Electron microscopy images of the wild type and *zip1Δ* were obtained from the figures of ^4^. The chromosome axis distance was estimated based on the size of the scale bar (Extended Data Fig. 7). Pixel size was calibrated using the scale bar on the image and the measurement was performed by Fiji.

### Statistical tests

Statistical tests were performed with R.

### Code availability

Custom code for low SNR microscopy system is available from the link below (https://github.com/frdchang/fcMatlabTools) ^42^.

