## Extended Data Fig. 1-9, Supplementary Table 1 for "Rapid Homolog Juxtaposition During Meiotic Chromosome Pairing"

### Extended Data Figures

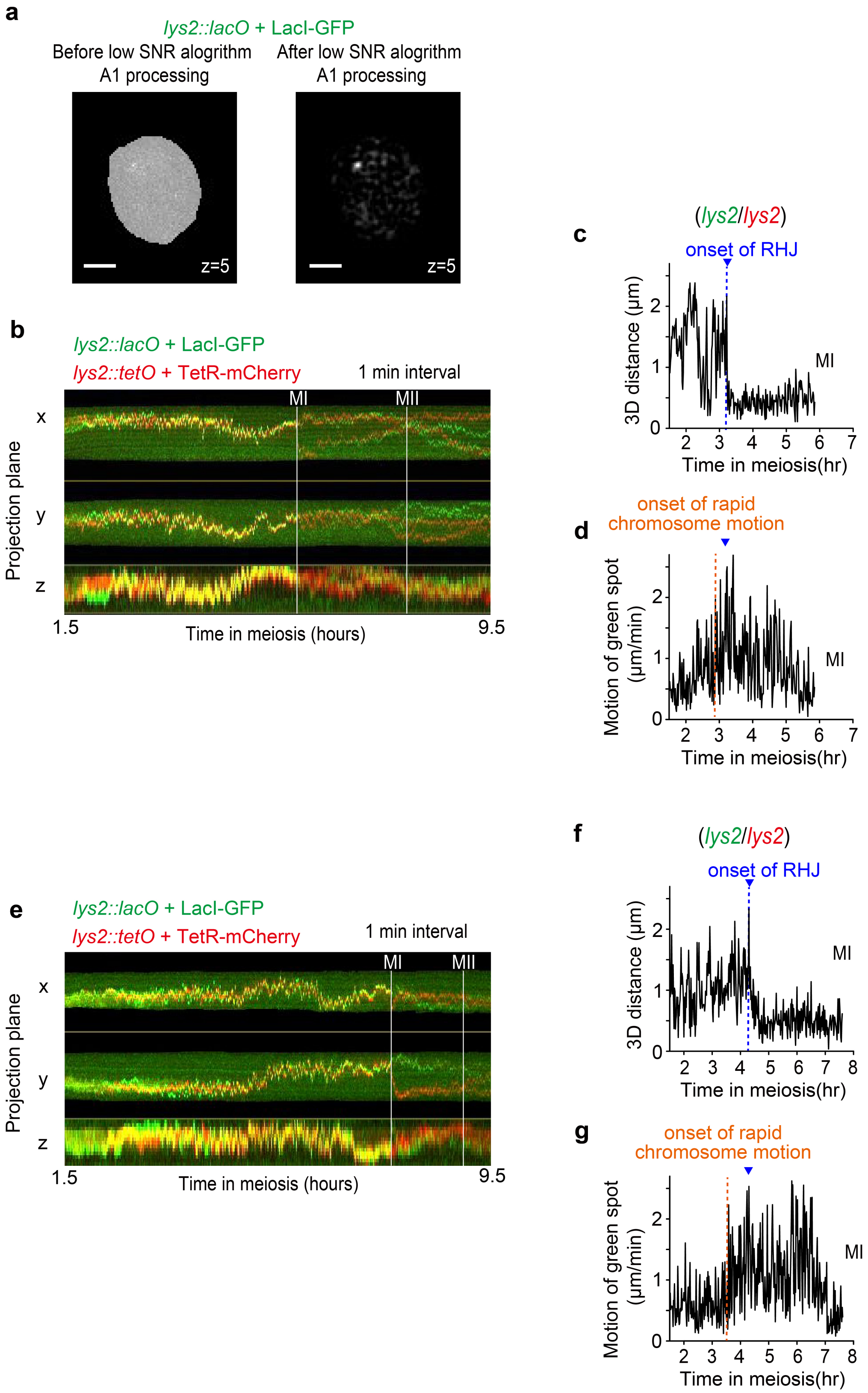


**Extended Data Fig. 1**: Long-timescale imaging of homologous loci during meiosis. **a**. The example of image processing of low SNR microscopy systems. The left figure is the raw data (after cell segmentation), and the right figure is the A1 filter processed image at the same Z-position. The scale bar is 2 µm. **b-g**. Trajectories of homolog loci through meiosis. **b**, **e**. Two representative kymographs of *lys2* homolog loci based on the imaging data with 1-minute intervals. The meiotic I division and meiotic II division time points are indicated by the white lines. **c**, **f**. The 3D distance plot between homologous loci through meiosis until MI division. **d**, **g**. The motion of the green spot in 1 min was plotted. The orange line indicates the time point of the onset of rapid chromosome motion. The blue triangle indicates the timing of the onset of RHJ.

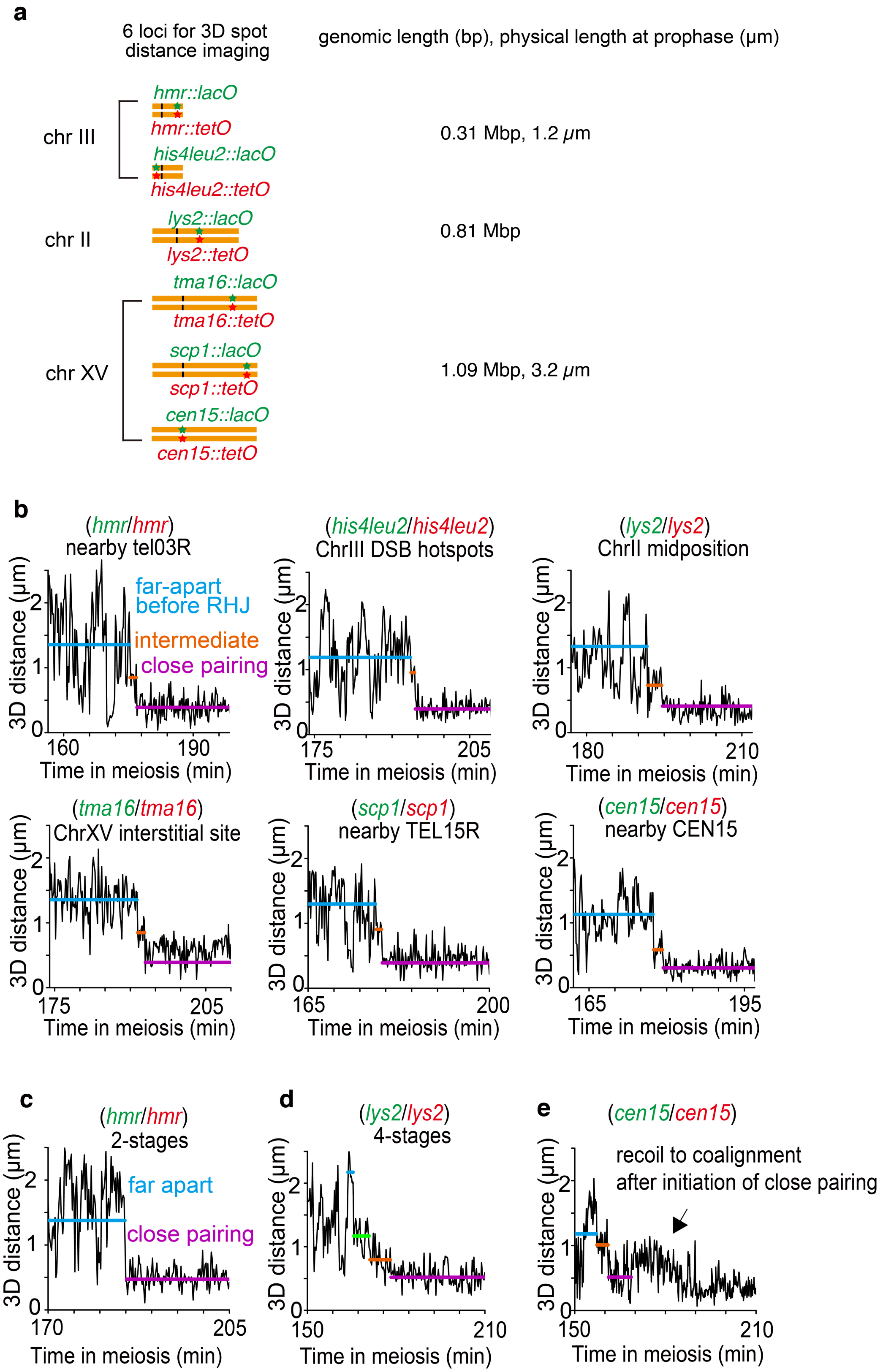


**Extended Data Fig. 2**: RHJ is a general process in meiotic budding yeast. **a**. The list of the fluorescent spot labeling used in this study with genomic and physical length. The physical length was estimated from the imaging of spread chromosomes in our previous paper [36]. The data of 3D spot distances from 60 cells were derived from 6 locus pairs: *lys2-lys2* (n=14 cells), *hmr-hmr* (n=10 cells), *tma16-tma16* (n=11 cells), *his4leu2-his4leu2* (n=8 cells), *scp1-scp1* (n=9 cells), and *cen15-cen15* (n=8 cells). **b**. The representative 3D distance plots exhibit RHJ at each locus. Each line indicates the average value at the far-apart stage (before RHJ, cyan), intermediate stage (orange), and close pairing stage (purple). **c**. The example of RHJ skipping the intermediate stage (2 stages and single step RHJ). This type of RHJ was quite rare in the wild type (4/61 cells, 6 locus pairs). **d**. The example of RHJ with an extra step is shown by a green line (4 stages and 4 stage RHJ). This cell was excluded from the general analysis in Fig. 2 (data of n=60 cells). However, this cell was included in the analysis to compare to the *ndj1Δ and csm4Δ* mutant cells and LatB-treated cells in Fig. 5. Such an extra step is rarely observed in the wild type cells (1/61 cells, 6 locus pairs). **e**. The example of RHJ with temporary recoiling to the intermediate after the close pairing. Cells with the homologous loci close to CenXV sometimes showed this pattern. However, this pattern could not be observed in other strains (5/61 cells, 6 locus pair but all of five cells were *cen15* loci).

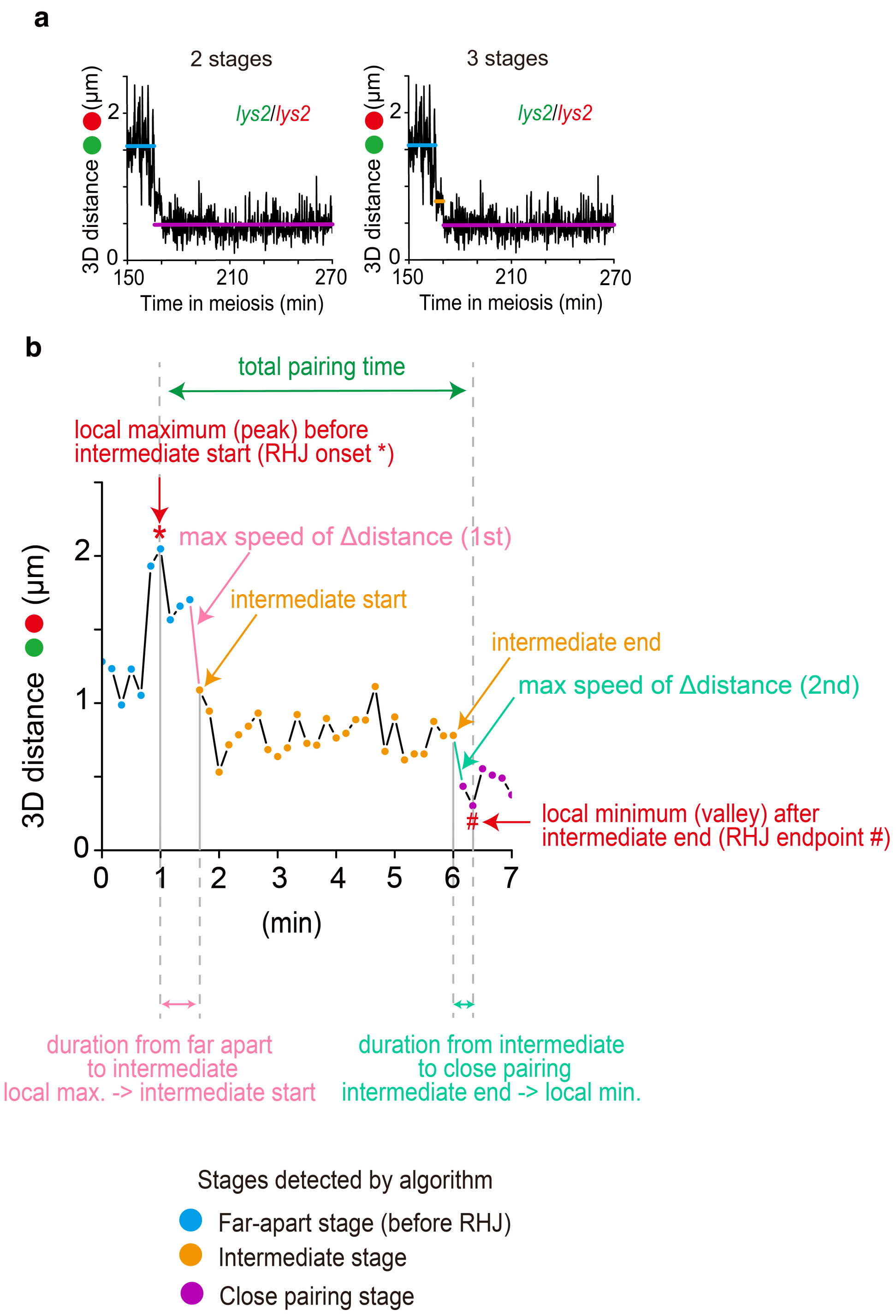


**Extended Data Fig. 3**: Analysis process of 3D distance changes. **a**. The step detection process with 2 stages (left) and 3 stages (right). **b**. Stages detected by the algorithm, RHJ onset/end, total pairing time, max speed of distance change, and transitions were summarized. The detailed process to detect the onset timing of the RHJ (local maximum, peak) and RHJ endpoint (local minimum, valley) was described in Materials and Methods (local maximum/minimum search). The 3D distance plot was created with a slight modification of the real data to explain the analysis process briefly.

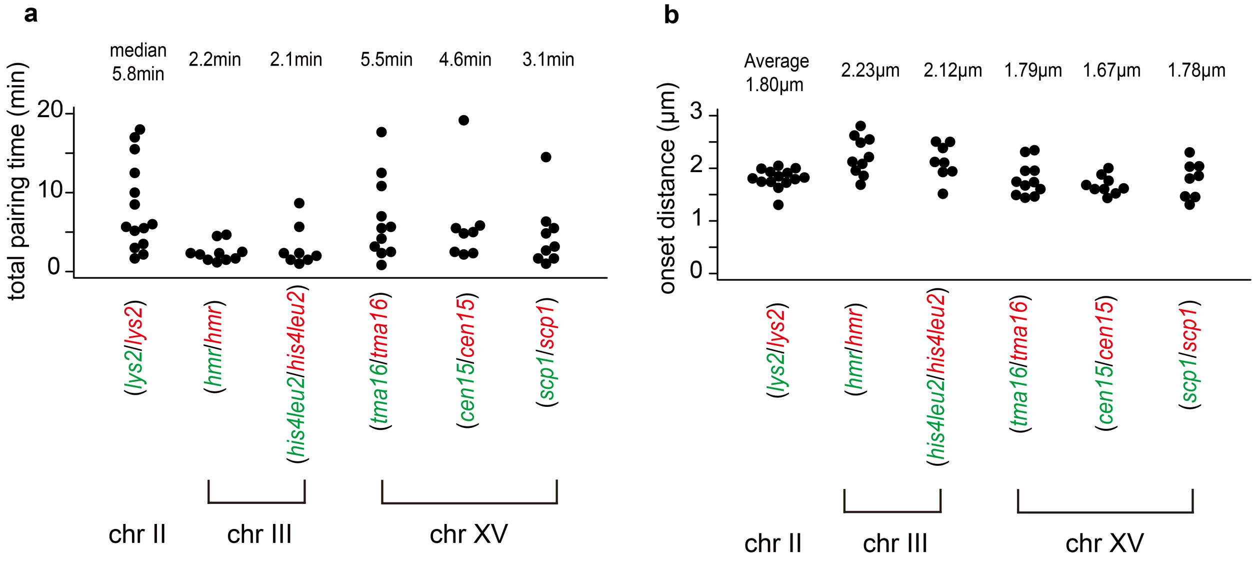


**Extended Data Fig. 4**: Total pairing time and RHJ onset distance for each locus. **a**, **b**. Distributions of the total pairing time (**a**) and the RHJ onset distance (**b**) for each locus are shown with median values and average values among cells, respectively.

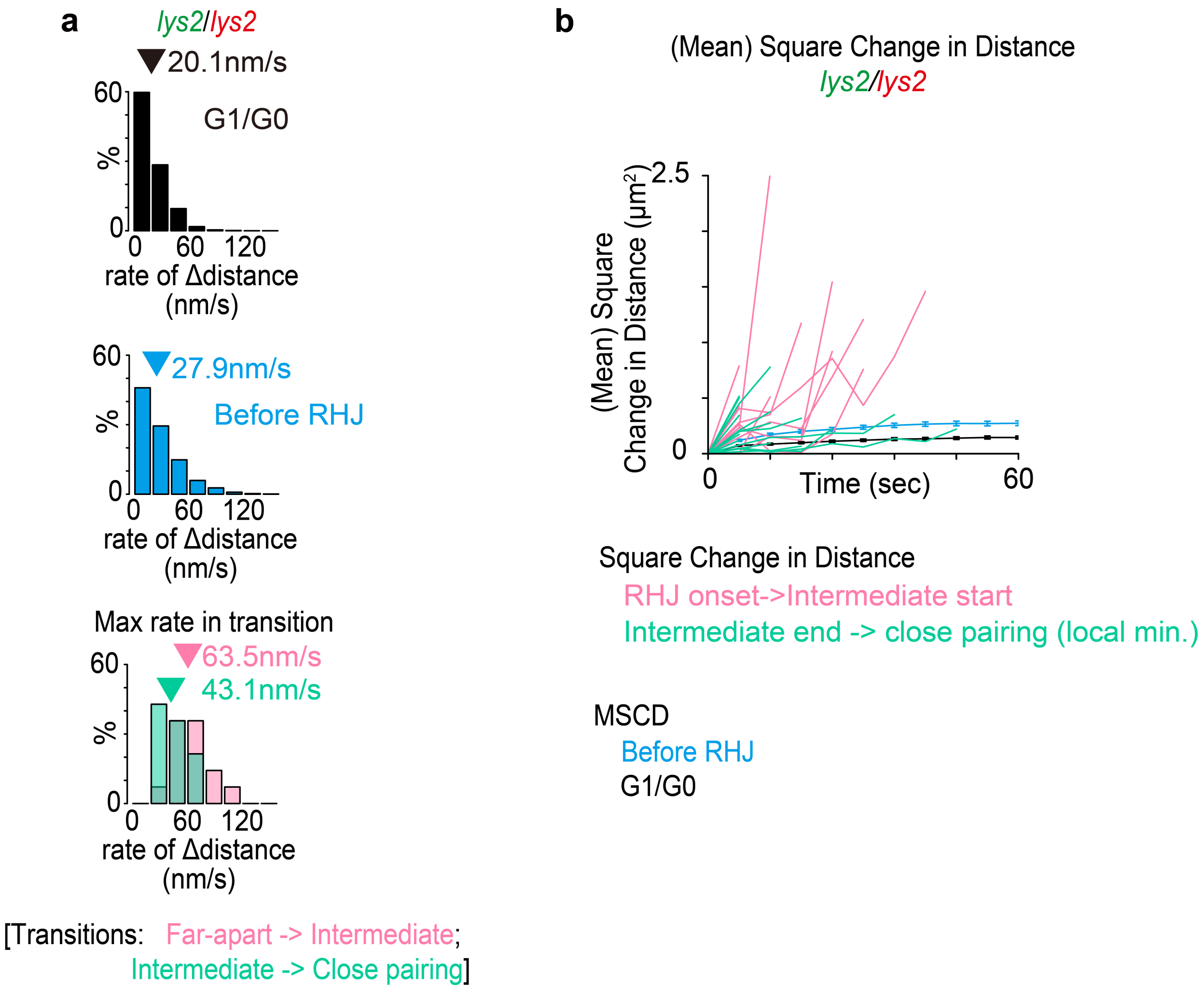


**Extended Data Fig. 5**: The detail of distance change analysis of *lys2* homologous loci. **a**. The distribution of distance change of homologous loci at G1/G0 (top), before RHJ (middle) and the maximum change in distances between homologous loci at the transitions from the far-apart to the intermediate (pink) and from the intermediate to the close pairing (light green). Triangles indicate the average. **b**. The lines of mean square change in distance (MSCD) at G1/G0 (black) and before RHJ (cyan) were shown with 95 % confidence intervals. The lines of square change in distance for each trajectory from the RHJ onset to the start of intermediate (pink) and from the end of intermediate to close pairing (light green) were also plotted. All lines of square change in distance from the RHJ onset to the start of intermediate exceeded the MSCD for the time points before RHJ (14/14 trajectories, pink). Most of the lines of square change in distance from the end of intermediate to the close pairing exceeded the MSCD for the time points before RHJ (9/14 trajectories, light green). The analysis shown here strongly implies the active process underlying the RHJ. Every data shown here was based on *lys2* homologous loci.

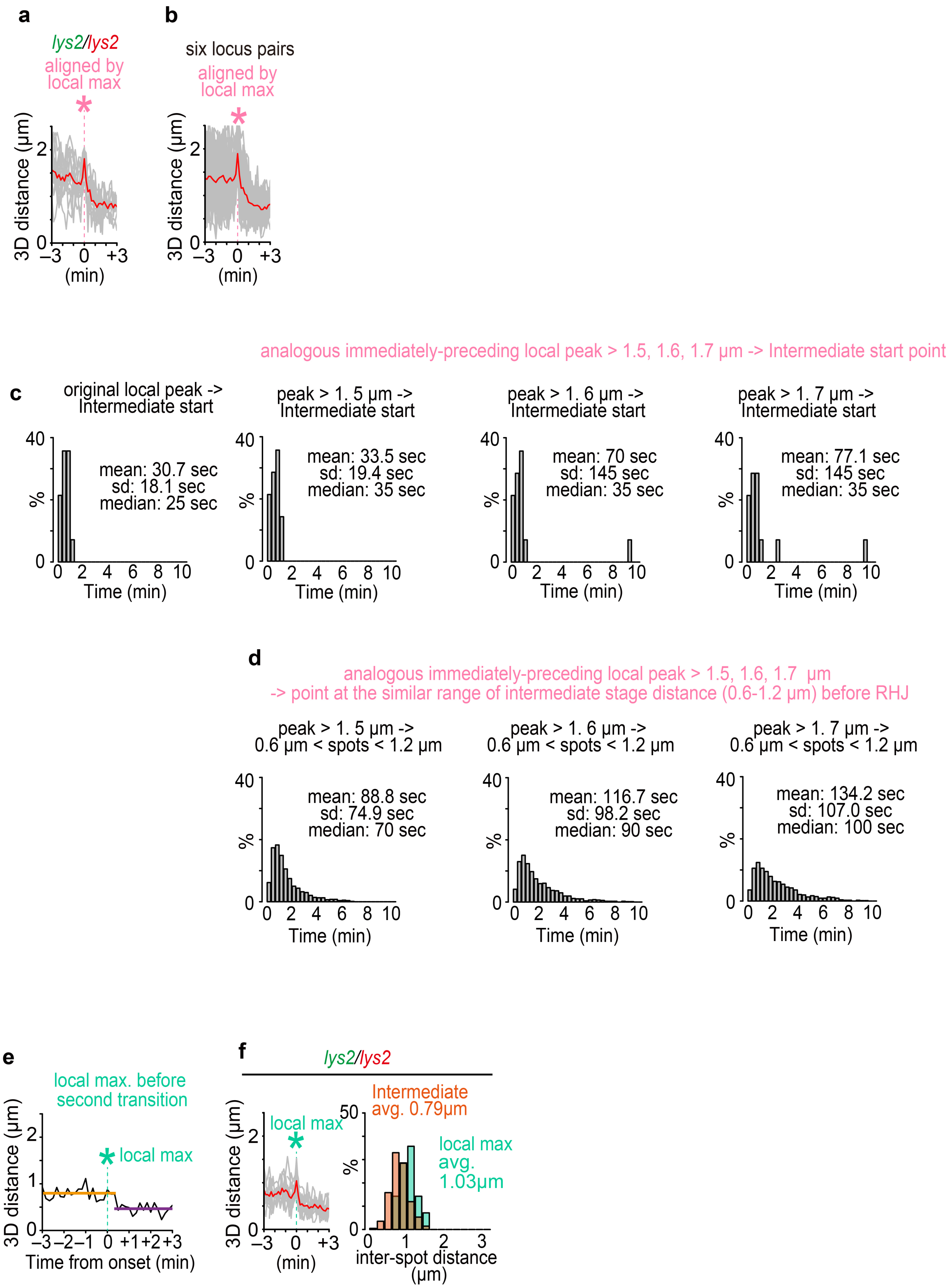


**Extended Data Fig. 6**: Time difference between the analogous immediately-preceding peak and the intermediate start. **a**, **b**. The plots of 3D distances between homolog loci for *lys2* (**a**) and six locus pairs (**b**) are aligned by the onset timing of RHJ (local maximum) (n=14 and 60 cells, respectively). The red line indicates the average of the cells. **c**, **d**. The time difference between the intermediate stage and immediately-preceding peak (local maximum, backward) or original local peak (**c**), and the time difference between the time point before RHJ whose distance is in the range of the intermediate stage (>0.6 µm and <1.2 µm) and the analogous immediately-preceding peak. The figures of **c** left and **d** left are the same as Fig. 2k pink and cyan figures. Several thresholds (1.5, 1.6, and 1.7 µm) were used to examine the relationship between the height of the analogous immediately-preceding peak and the intermediate start point (and the control point whose distance is in the same range as the intermediate stage). However, the result was not different in each condition, suggesting that RHJ onset is triggered by extension along a nascent inter-homolog linkage. **e**. The example of the transition from intermediate to close pairing (*lys2* homolog loci). The transition onset is defined by the local peak before the start of close pairing. **f**. (left) The distance plots among 14 cells (*lys2* homolog loci) are aligned by the local maximum before the 2nd transition (intermediate to close pairing). The red line indicates the average of the cells. (right) Distance distribution of the intermediate stage (orange) and local maximum before the 2nd transition (light green).

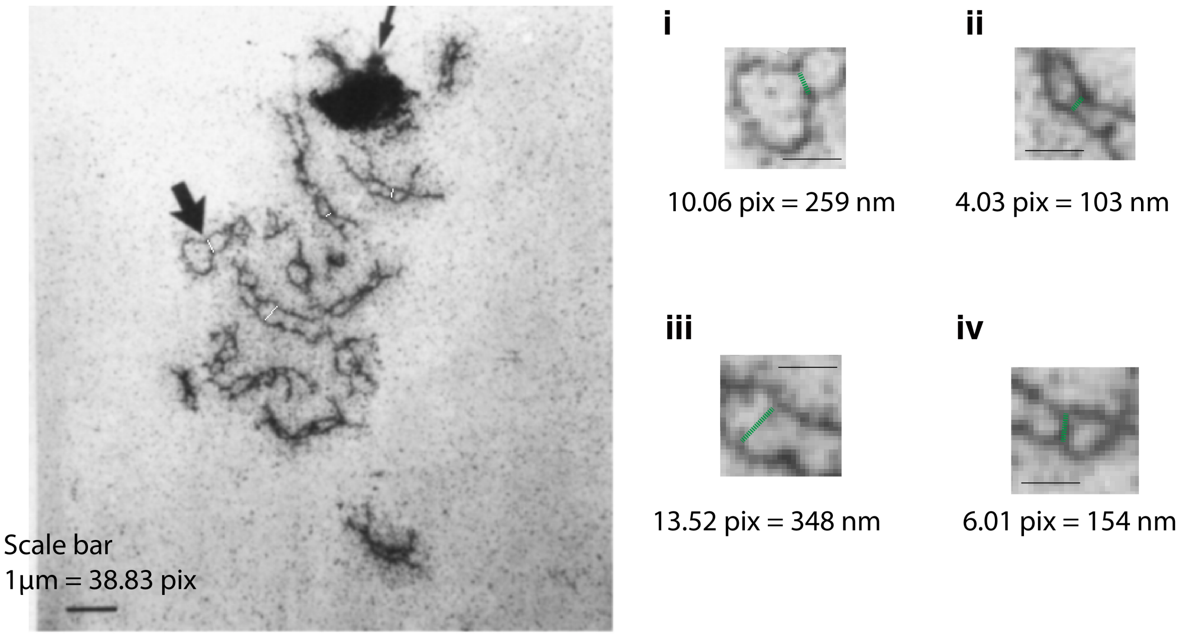


**Extended Data Fig. 7**: Analysis of the distance between chromosome axes in electron microscopy image of *zip1Δ* mutant cell. Data was based on the paper of Sym et al. 1993 (ref 4). The white lines in the left figure and the green lines in the right figure are at the same position and these lines are used to measure the distance between chromosome axes. The scale bar in the left figure is 1 µm (original) and the scale bars in the right figure are 500 nm (added by this study).

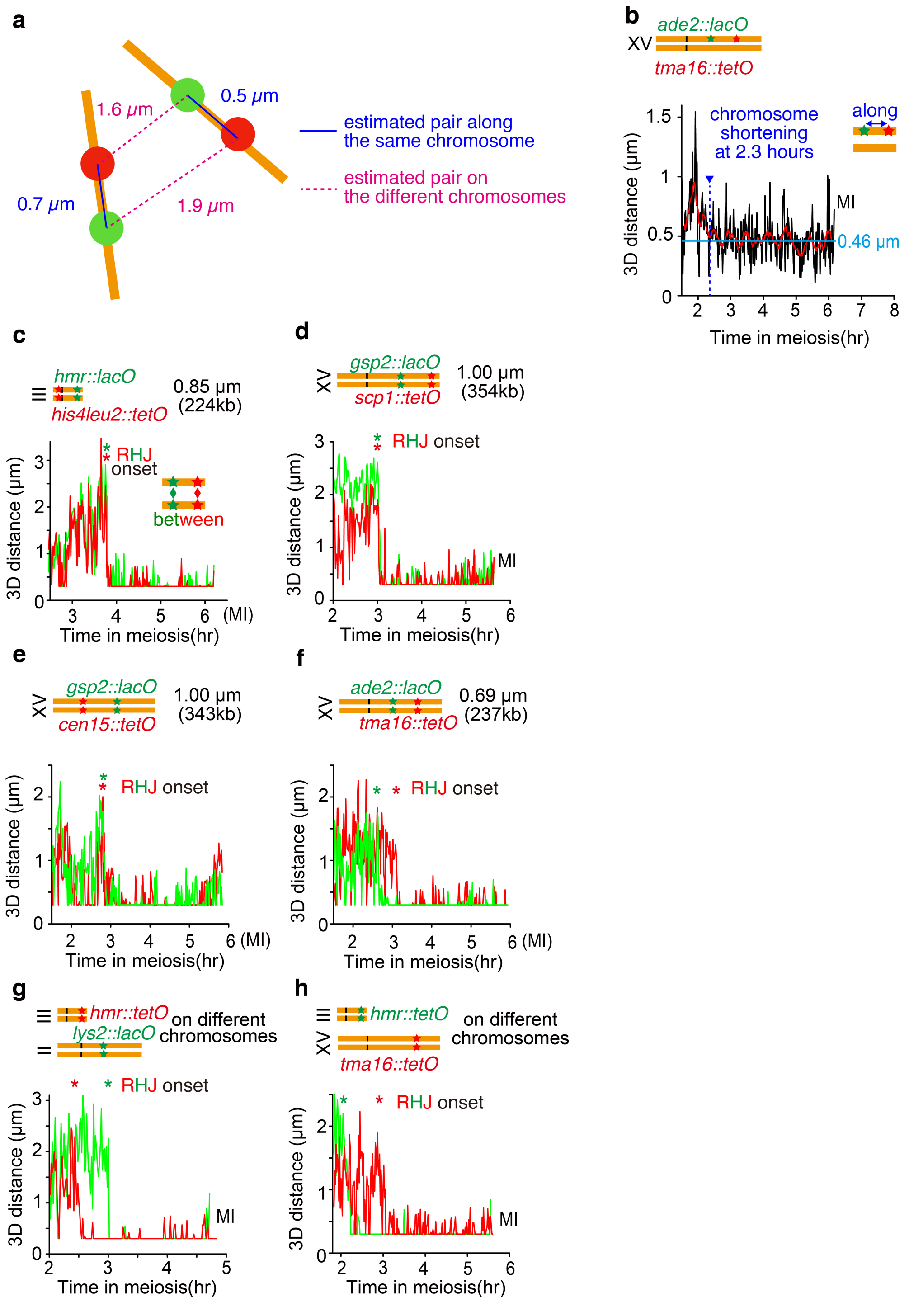


**Extended Data Fig. 8**: Multiple spots imaging for RHJ onset timing and chromosome shortening. **a**. The procedure of the longitudinal distance analysis for fluorescent loci along the chromosomes. There are two combinations of green and red spots (blue lines and pink dot lines) when every four spots are separately observed. The combination of the spots along the same chromosomes is estimated by the minimalization of the maximum distance between spots (e.g., 1.9 μm vs 0.7 μm -> 0.7 μm in the figure). **b**. The longitudinal spots analysis on the single chromosome labeling. The red line is the average in 10 min. The chromosome shortening timing is indicated by a blue dotted line. The average distance after the chromosome shortening was 0.46 μm and plotted by cyan line. After the chromosome shortening, the distance between spots along the chromosome was maintained stably by the MI division with a small fluctuation. **c-h**. The representative examples of pairing at multiple spots along the chromosomes. **c**. *hmr-his4leu2* pairing along chromosome IIIs. **d**. *gsp2-scp1* pairing along chromosome XVs. **e**. *gsp2-cen15* pairing along chromosome XVs. **f**. The unsynchronized case of *ade2-tma16* pairing. **g, h**. The unsynchronized cases of pairing at different chromosomes.

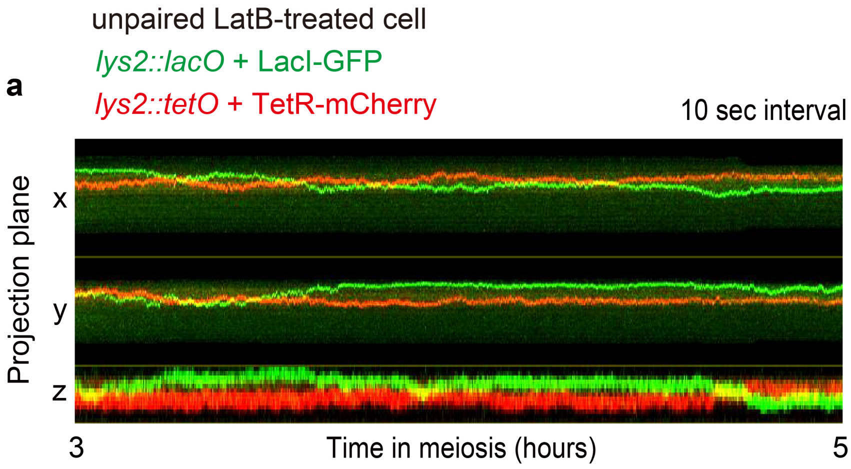


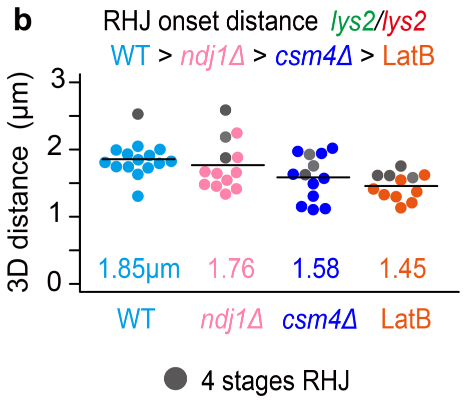


**Extended Data Fig. 9**: The details of the RHJ in rapid chromosome motion mutant analysis. **a**. The representative example of unpaired *lys2* homologous loci by 5 hours in meiosis in a LatB-treated cell. ~50% of the cells were shown this pattern in LatB-treated cells. **b**. RHJ onset distance distribution between *lys2* loci in wild type cells, *ndj1Δ* and *csm4Δ* mutant cells, and LatB-treated cells with the indication of 4 stage RHJ (gray circles).

**Supplementary Video 1**: The long-timescale imaging of *lys2* homologous loci labeled by *lacO* array/LacI-mEGFP (green) and *tetO* array/TetR-mCherry (red) at meiotic prophase (2 hours in SPM to MI/MII divisions). The 3D-stack images were taken once per 1 minute.

**Supplementary Video 2**: The short-timescale imaging of *lys2* homologous loci labeled by *lacO* array/LacI-mEGFP (green) and *tetO* array/TetR-mCherry (red) at meiotic prophase (2.5 - 4.5 hours in SPM). The 3D-stack images were taken once per 10 seconds.

**Supplementary Video 3**: The short-timescale imaging of *hmr* homologous loci labeled by *lacO* array/LacI-mEGFP (green) and *tetO* array/TetR-mCherry (red) at meiotic prophase (2.5 - 4.5 hours in SPM). The 3D-stack images were taken once per 10 seconds.

**Supplementary Video 4**: The short-timescale imaging of *lys2* homologous loci labeled by *lacO* array/LacI-mEGFP (green) and *tetO* array/TetR-mCherry (red) at meiotic pachytene stage in the *ndt80Δ* mutant cell (8 – 8.5 hours in SPM). The 3D-stack images were taken once per 10 seconds.

**Supplementary Video 5**: The long-timescale imaging of *ade2-tma16* homologous loci labeled by *lacO* array/LacI-mEGFP (green) and *tetO* array/TetR-mCherry (red) at meiotic prophase (2 hours in SPM to MI division). The 3D-stack images were taken once per 1 minute.

**Supplementary Video 6**: The long-timescale imaging of *hmr-his4leu2* homologous loci labeled by *lacO* array/LacI-mEGFP (green) and *tetO* array/TetR-mCherry (red) at meiotic prophase (2 hours in SPM to MI division). The 3D-stack images were taken once per 1 minute.

**Supplementary Video 7**: The long-timescale imaging of *lys2/lys2* and *hmr/hmr* homologous loci labeled by *lacO* array/LacI-mEGFP (green) and *tetO* array/TetR-mCherry (red) at meiotic prophase (1.5 hours in SPM to MI division). The 3D-stack images were taken once per 1 minute.

**Supplementary Video 8**: The short-timescale imaging of *lys2* homologous loci labeled by *lacO* array/LacI-mEGFP (green) and *tetO* array/TetR-mCherry (red) at meiotic prophase in the *ndj1* mutant cell (4 - 5 hours in SPM). The 3D-stack images were taken once per 10 seconds.

**Supplementary Table 1**: The list of budding yeast strains used in this study.

All strains are MATa/MATα derivatives of SK1 background.

| Stain | Genotype |
|---|---|
| NKY4265 | *ho::hisG/", lys2::TetO array::URA3/lys::LacO array::URA3, leu2::tetR-tdmCherry-LEU2/leu2::hisG, ura3::LacI-mEGFP-URA3/ura3 (ΔPstI-SmaI), nuc1::CUP1Promoter-LacI-mEGFP::KanMX/NUC1* |
| NKY4266 | *ho::hisG/", hmr::TetO array::URA3/hmr::LacO array::URA3, leu2::tetR-tdmCherry-LEU2/leu2::hisG, ura3::LacI-mEGFP-URA3/ura3 (ΔPstI-SmaI), nuc1::CUP1Promoter-LacI-mEGFP::KanMX/NUC1* |
| NKY4267 | *ho::hisG/", HIS4::LEU2(BamHI)::LacO array/his4X::LEU2 (NgoMIV)::TetO array, leu2::tetR-tdmCherry-LEU2/", ura3::LacI-mEGFP-URA3/ura3 (ΔPstI-SmaI), nuc1::CUP1Promoter-LacI-mEGFP::KanMX/"* |
| NKY4268 | *ho::hisG/", tma16::LacO array::URA3/tma16::TetO array::URA3, leu2::tetR-tdmCherry-LEU2/leu2::hisG, ura3::LacI-mEGFP-URA3/ura3 (ΔPstI-SmaI), nuc1::CUP1Promoter-LacI-mEGFP::NAT/nuc1::CUP1Promoter-LacI-mEGFP::KanMX* |
| NKY4269 | *ho::hisG/", scp1::LacO array::LEU2/scp1::TetO array::LEU2, leu2::tetR-tdmCherry-LEU2/leu2::hisG, ura3::LacI-mEGFP-URA3/ura3(ΔPstI-SmaI), nuc1::CUP1Promoter-LacI-mEGFP::KanMX/NUC1* |
| NKY4270 | *ho::hisG/", cen15::LacO array::KanMX/cen15::TetO array::KanMX, leu2::tetR-tdmCherry-LEU2/leu2::hisG, ura3::LacI-mEGFP-URA3/ura3(ΔPstI-SmaI)::hisG, nuc1::CUP1Promoter-LacI-mEGFP::NAT/NUC1,* |
| NKY4271 | *ho::hisG/", spo11Y135F::Hyg/", lys2::LacO array::URA3/lys2::TetO array::URA3, leu2::tetR-tdmCherry-LEU2/leu2::hisG, ura3::LacI-mEGFP-URA3/ura3 (ΔPstI-SmaI), nuc1::CUP1Promoter-LacI-mEGFP::KanMX/NUC* |
| NKY4272 | *ho::hisG/", dmc1::KanMX/dmc1::LEU2, lys2::LacO array::URA3/lys2::TetO array::URA3, leu2::tetR-tdmCherry-LEU2/leu2::hisG,ura3::LacI-mEGFP-URA3/", nuc1::CUP1Promoter-LacI-mEGFP::KanMX/NUC1* |
| NKY4273 | *ho::hisG/", hop2::KanMX/", lys2::LacO array::URA3/lys2::TetO array::URA3, leu2::tetR-tdmCherry-LEU2/leu2::hisG, ura3::LacI-mEGFP-URA3/ura3 (ΔPstI-SmaI), nuc1::CUP1Promoter-LacI-mEGFP::NAT/NUC1* |
| NKY4274 | *ho::hisG/", ndt80::LEU2/", lys2::LacO array::URA3/lys2::TetO array::URA3, leu2::tetR-tdmCherry-LEU2/leu2::hisG, ura3::LacI-mEGFP-URA3/ura3(ΔPstI-SmaI)* |
| NKY4275 | *ho::hisG/", zip1::KanMX/zip1::LEU2, lys2::LacO array::URA3/lys2:TetO array::URA3, leu2::tetR-tdmCherry-LEU2/leu2::hisG, ura3::LacI-mEGFP-URA3/ura3 (ΔPstI-SmaI)* |
| NKY4276 | *ho::hisG/", syc1::LacO array::NAT/", tma16::TetO array::URA3/", leu2::tetR-tdmCherry-LEU2/leu2::hisG, ura3::LacI-mEGFP-URA/", nuc1::CUP1Promoter-LacI-mEGFP::KanMX/"* |
| NKY4277 | *ho::hisG/", hmr::LacO array::URA3/", his4X::LEU2 (NgoMIV)::TetO array/", leu2::tetR-tdmCherry-LEU2/leu2::hisG, ura3::LacI-mEGFP-URA/", nuc1::CUP1Promoter-LacI-mEGFP::KanMX/NUC1* |
| NKY4278 | *ho::hisG/", ade2::LacO array-KanMX/", tma16::TetO array::URA3/", leu2::tetR-tdmCherry-LEU2/", ura3::LacI-mEGFP-URA3/", nuc1::CUP1Promoter-LacI-mEGFP::NAT/", his4X::ADE2-his4B/HIS4* |
| NKY4279 | *ho::hisG/", gsp2::LacO array::hphMx4/", cen15::TetO array::KanMX4/", leu2::tetR-tdmCherry-LEU2/", ura3::LacI-mEGFP-URA/", nuc1::CUP1Promoter-LacI-mEGFP::KanMX/NUC1* |

| | |
|---|---|
| NKY4280 | *ho::hisG/", gsp2::LacO array::hphMx4/", scp1::TetO array::LEU2/", leu2::tetR-mCherry-LEU2/leu2::tetR-tdmCherry-LEU2, ura3::LacI-mEGFP-URA/", nuc1::CUP1Promoter-LacI-mEGFP::KanMX/NUC1* |
| NKY4281 | *ho::hisG/", lys2::LacO array::URA3/", hmr::TetO array::URA3/", leu2::tetR-tdmCherry-LEU2/", ura3::LacI-mEGFP-URA3/ura3 (ΔPstI-SmaI), nuc1::CUP1Promoter-LacI-mEGFP::NAT/NUC1* |
| NKY4282 | *ho::hisG/", hmr::LacO array::URA3/", tma16::TetO array::URA3/", leu2::tetR-tdmCherry-LEU2/", ura3::LacI-mEGFP-URA/", nuc1::CUP1Promoter-LacI-mEGFP::NAT/"* |
| NKY4283 | *ho::hisG/", tma16::TetO array::URA3/TMA16, ade2::LacO array::KanMX/ADE2, leu2::tetR-tdmCherry-LEU2/", ura3::LacI-mEGFP-URA/", nuc1::CUP1Promoter-LacI-mEGFP::NAT/"* |
| NKY4284 | *ho::hisG/", ndj1::Hyg/", lys2::LacO array::URA3/lys2::TetO array::URA3, leu2::tetR-tdmCherry-LEU2/leu2::hisG, ura3::LacI-mEGFP-URA3/ura3 (ΔPstI-SmaI)* |
| NKY4285 | *ho::hisG/", csm4::Hyg/", lys2::LacO array::URA3/lys2::TetO array::URA3, leu2::tetR-tdmCherry-LEU2/", ura3::LacI-mEGFP-URA3/ura3 (ΔPstI-SmaI), nuc1::CUP1Promoter-LacI-mEGFP::NAT/NUC1* |
